# The limited spatial scale of dispersal in soil arthropods revealed with whole-community haplotype-level metabarcoding

**DOI:** 10.1101/2020.06.18.159012

**Authors:** P Arribas, C Andújar, A Salces-Castellano, BC Emerson, AP Vogler

**Affiliations:** Island Ecology and Evolution Research Group (IPNA-CSIC), Astrofísico Fco. Sánchez 3, 38206 La Laguna, Tenerife, Spain; Department of Life Sciences, Natural History Museum, Cromwell Road, London SW7 5BD, UK; Department of Life Sciences, Imperial College London, Silwood Park Campus, Ascot SL5 7PY, UK

**Keywords:** cMBC, dispersal, distance-decay, endemism, haplotype, soil mesofauna, speciation scale

## Abstract

Soil mesofauna communities are hyperdiverse and critical for ecosystem functioning. However, our knowledge on spatial structure and underlying processes of community assembly for soil arthropods is scarce, hampered by limited empirical data on species diversity and turnover. We implement a high-throughput-sequencing approach to generate comparative data for thousands of arthropods at three hierarchical levels: genetic, species and supra-specific lineages. A joint analysis of the spatial arrangement across these levels can reveal the predominant processes driving the variation in biological assemblages at the local scale. This multi-hierarchical approach was performed using haplotype-level-COI metabarcoding of entire communities of mites, springtails and beetles from three Iberian mountain regions. Tens of thousands of specimens were extracted from deep and superficial soil layers and produced comparative phylogeographic data for >1000 co-distributed species and nearly 3000 haplotypes. Local assemblages were highly distinctive between grasslands and forests, and within each of them showed strong spatial structures and high endemicity at the scale of a few kilometres or less. The local distance-decay patterns were self-similar for the haplotypes and higher hierarchical entities, and this fractal structure was very similar in all three regions, pointing to a significant role of dispersal limitation driving the local-scale community assembly. Our results from whole-community metabarcoding provide unprecedented insight into how dispersal limitations constrain mesofauna community structure within local spatial settings over evolutionary timescales. If generalized across wider areas, the high turnover and endemicity in the soil locally may indicate extremely high richness globally, challenging our current estimations of total arthropod-diversity on Earth.

## INTRODUCTION

Soils are among the most biodiverse habitats on Earth, but represent probably the least well studied, and thus poorly understood, terrestrial ecosystem (Bardgett & van der Putten, 2014; Decaëns, 2010). Current understanding of terrestrial biodiversity has mainly relied on studies of aboveground organisms, but in recent years efforts have been focussed on developing an integrative understanding of soil biodiversity patterns and underlying mechanisms (Thakur et al., 2019). However, current knowledge on soil biodiversity is strongly unbalanced across taxonomic groups (Cameron et al., 2018), which hampers the development of such integrative framework of soil biodiversity. In particular, there is a pronounced shortage of basic biodiversity data, including estimations of their species diversity and how it is spatially structured, for the taxonomically and functionally diverse soil arthropod mesofauna composed of small-bodied invertebrates measuring between 0.1 – 2 mm and found by thousands in virtually every square meter of natural soil (Bardgett, Usher, & Hopkins, 2005; Decaëns, 2010). In recent years, high-throughput sequencing has been applied widely to study the microbial components of soil ecosystems and led to a solid understanding of global distribution patterns and the ecological drivers of soil microbiomes (e.g. Delgado-Baquerizo et al., 2018; Ramirez et al., 2018, 2014). In contrast, the study of soil arthropod mesofauna has seen comparatively little progress in exploiting these tools (mostly using 18S eDNA approaches Wu, Ayres, Bardgett, Wall, & Garey, 2011; Zinger et al., 2019), whereas conventional taxonomic approaches have been onerous, given the small body size, limited morphological variation and high local abundances of most mesofauna components.

Existing work on the diversity, distribution and community composition of soil arthropods has focussed on springtails and oribatid mites, and mostly has pointed to selection by abiotic and/or biotic environmental factors as major mechanisms of community assembly at the local scale (e.g. Caruso, Trokhymets, Bargagli, & Convey, 2013; Magilton, Maraun, Emmerson, & Caruso, 2019; reviewed in Berg, 2012; Thakur et al., 2019). Different studies have also reported purely spatial structures (independent of the measured environmental variables) or stochastic patterns (non-environmental neither spatial structures) for the soil mesofauna communities, that have been recurrently attributed to the contribution of demographic processes (i.e., ecological drift without dispersal limitation) in determining the local community assembly (Bahram, Kohout, Anslan, Harend, & Abarenkov, 2016; Ingimarsdóttir et al., 2012; Widenfalk, Malmström, Berg, & Bengtsson, 2016; Zinger et al., 2019). In addition, dispersal limitations have also been suggested to contribute to some of the spatial community structures reported (Caruso, Taormina, & Migliorini, 2012; Gao, He, Zhang, Liu, & Wu, 2014), but still is rarely recognised as an important mechanism of assembly of the soil mesofauna at the local scale (Berg, 2012; Thakur et al., 2019). Beyond the aggregated distribution within the geographic ranges of the species, limitations to dispersal can determine the degree to which species pools are differentiated over spatial distance (Hortal, Roura-Pascual, Sanders, & Rahbek, 2010). These effects are frequently evident as biogeographic or phylogeographic breaks at large (regional to continent-wide) scales, but if the scale of movement is highly constrained and if the constraints are persistent through time, as could be the case in the soil matrix, purely spatial patterns of community assemblage dominated by species (and haplotype) turnover can arise even over relatively small distances. The potentially low taxonomic resolution (due to morphological species assignment or the use of 18S rRNA gene) of most of the studies on arthropod mesofauna communities may have missed the importance of dispersal limitation in determining the diversity patterns of soil mesofauna (but see Andújar et al., 2015; Lindo & Winchester, 2009).

The spatial scale at which dispersal constraints are effective in determining species distributions and community assembly is a major “open question” in soil biodiversity research (Thakur et al., 2019). For the mesofauna, small body size and high local abundance may increase the probability of passive dispersal and long-distance movement, and therefore dispersal constraints within the soil may be of limited importance. In fact, a high prevalence of aerial, aquatic and marine rafting has been demonstrated for various mesofaunal lineages (Coulson, Hodkinson, Webb, & Harrison, 2002; Nkem et al., 2006; Schuppenhauer, Lehmitz, & Xylander, 2019), and studies have shown mesofaunal assemblages with no apparent dispersal limitation across continental-scale areas (Baird, Leihy, Scheepers, & Chown, 2019), especially for the smallest-bodied soil arthropods (Gan, Zak, & Hunter, 2019). On the other hand, molecular studies have revealed high differentiation and ancient microendemicity even in morphologically indistinguishable clades, indicating long-term constraints to dispersal (Andújar, Pérez-González, et al., 2017; Cicconardi, Fanciulli, & Emerson, 2013). These empirical data limited to particular mesofauna lineages and their contrasting findings highlight the difficulty of establishing the role of dispersal constraints in community assembly. As such, inferences regarding the distribution and diversification of edaphic species, and thus generalisations regarding macroecological and macroevolutionary patterns, remain challenging.

New approaches to the study of diverse and cryptic arthropods using whole-community metabarcoding (cMBC) using the mitochondrial COI gene are now revolutionizing the understanding of complex arthropod communities (Arribas et al., 2016; Ji et al., 2013). The methodology involves the bulk sequencing of mixed communities and subsequent clustering of DNA reads into operational taxonomic units (OTUs) that broadly represent the species category. While an efficient method to approximate community profiles at the species-level, precise removal of primary DNA reads affected by sequencing errors (Andújar, Arribas, Yu, Vogler, & Emerson, 2018; Elbrecht, Vamos, Steinke, & Leese, 2018; Turon, Antich, Palacín, Præbel, & Wangensteen, 2019) and co-amplified nuclear mitochondrial copies (numts) (Andújar et al. under review) obviate the need for clustering. Read-based data raise the prospect of reliable haplotype information from mitochondrial COI cMBC, which represents a step change for the study of diversity patterns through whole-community genetic analyses at haplotype-level resolution.

The availability of metabarcode data at both species and haplotype levels permits the joint analysis of turnover (beta diversity) at different hierarchical levels, to assess whether the variation in biological assemblages is predominantly driven by dispersal or niche-based processes (Baselga et al., 2013; Baselga, Gómez-Rodríguez, & Vogler, 2015). Local assemblages may diverge simply due to the lack of population movement which, when assessed for entire communities, results in a largely regular decay of community similarity with spatial distance for the typically neutral haplotype variation of the mitochondrial COI gene. Under a scenario where dispersal constraints determine the spatial community structure, assemblage turnover at the species level should mirror these haplotype patterns, albeit at a higher level of similarity. In contrast, niche-based processes acting on species traits produce species distributions that mainly follow environmental factors and thus differ from neutral conditions determining the haplotype distributions. This confounds the correlation (self-similarity) of distance decay at the species and haplotype levels, each driven by different processes, and thus the joint analysis of communities at genetic and species levels provides a formal test to discern if a particular spatial pattern of community assemblage is predominantly driven by dispersal (i.e., neutral) or niche-based processes (Baselga et al., 2013, 2015). In addition, multi-hierarchical analyses may also describe the spatial scale at which dispersal constraints act, and the variation of scale among different taxonomic groups or habitats (Gómez-Rodríguez, Miller, Castillejo, Iglesias-Piñeiro, & Baselga, 2018; Múrria et al., 2017). This framework remains to be exploited with whole-community mitochondrial cMBC.

Here we apply the multi-hierarchical framework to study the spatial structure of entire assemblages of mites (Acari), springtails (Collembola) and beetles (Coleoptera) including many thousands of specimens, in a semi-natural mosaic landscape within three geographically distinct mountain regions in southern and central Iberia (Fig. 1 A). Our aim was to generate rigorous whole-community data at haplotype, species (OTU) and supra-specific levels to evaluate the spatial turnover at the local scale (i.e. <10 km, following scale definitions of Pearson & Dawson, 2003) and in two habitat types within the same spatial settings. Using the three regions as natural replicates, we evaluated patterns of richness, endemicity, turnover and the spatial scale of the distance decay in community similarity at each hierarchical level and assessed the prevailing ecological and evolutionary processes that determine the diversity and spatial distribution of soil arthropod communities at the local scale.

**Figure 1.**
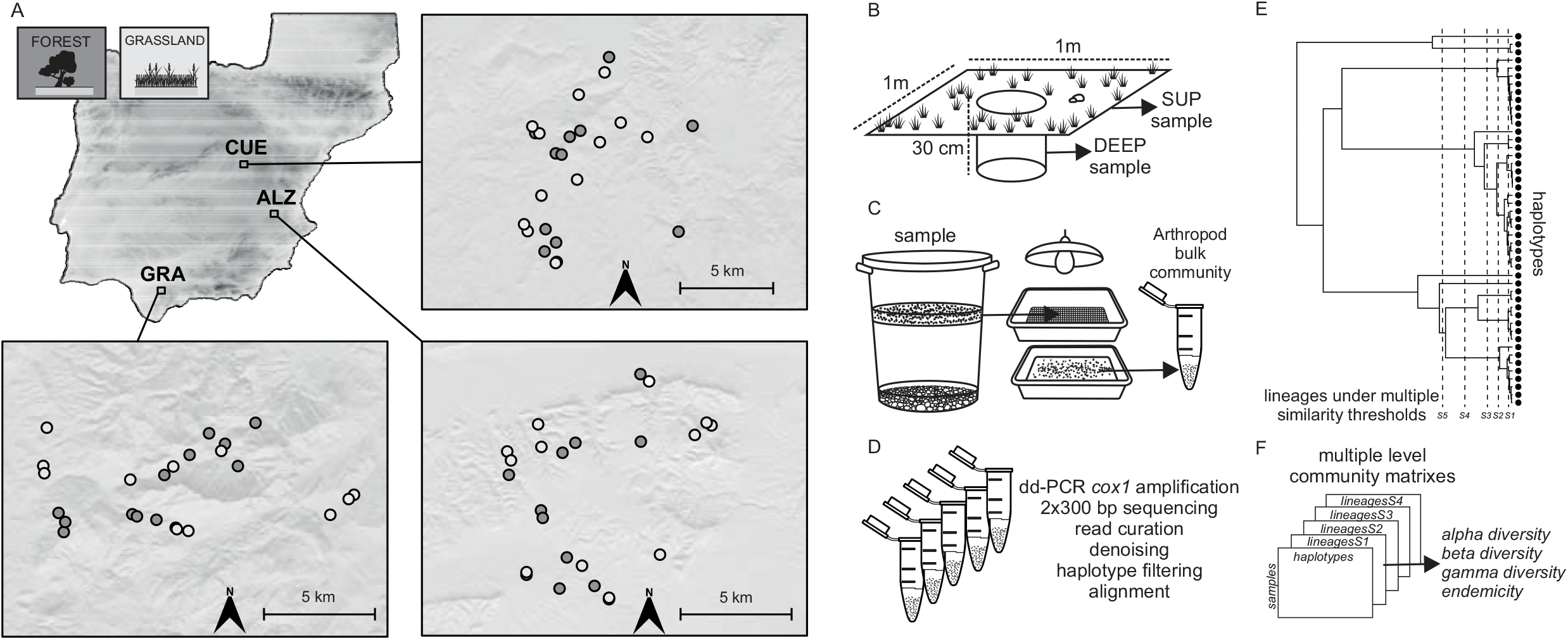
Sampling points in the three local settings within the Iberian Peninsula, Sierra de Grazalema, (GRA), Sierra de Alatoz (ALZ) and Sierra de la Alcarria Conquense (CUE). Sampling points are located within *Quercus* forest patches (dark grey) and wet grassland patches (pale grey).

## MATERIALS AND METHODS

### Soil sampling and mesofauna extraction

A total of 144 soil samples were collected from three regions in the southern Iberian Peninsula at Sierra de Grazalema, (GRA), Sierra de Alatoz (ALZ) and Sierra de la Alcarria Conquense (CUE) (Fig. 1 A). In each region, 24 points were sampled, half of them in *Quercus* forest and half in wet grassland habitat, at distances of 500 m to a maximum of 15 km (Fig. 1, Table S1). For each point, we collected i) a sample containing the superficial soil layer (SUP), by extracting one square meter of leaf litter and humus up to 5 cm deep and ii) a sample of the corresponding deep soil layer (DEEP), by extracting the substrate of a 30 cm diameter core to 30 cm depth, comprising ca. 20 litres of soil. Within each region and habitat, sampling points were located in natural patches of similar dominant vegetation and elevation. Different variables characterising the sampling points were recorded including elevation, slope, orientation, stoniness, humus depth, qualitative porosity, roots, soil temperature and soil relative humidity (Table S1).

Superficial and deep soil samples were processed following the flotation– Berlese–flotation protocol (FBF) of Arribas et al. (2016) for the ‘clean’ extraction of arthropod mesofauna from a large volume of soil. Briefly, the FBF protocol is based on the flotation of soil in water, which allows the extraction of the organic (floating) matter containing the soil mesofauna from raw soil samples. Subsequently, the organic portion is placed in a modified Berlese apparatus to capture specimens alive and preserve them in absolute ethanol. The last part of the FBF protocol includes additional flotation and filtering steps of the ethanol-preserved arthropods using 1-mm and 0.45-µm wire mesh sieves to remove debris and dirt accumulated in the Berlese extract. This procedure generates two ‘clean’ subsamples of bulk specimens for DNA extraction, one including all adult and larval Coleoptera, and a second with the smallest mesofauna typically dominated by mites and springtails.

### DNA extraction, PCR amplification and Illumina sequencing

Each bulk specimen subsample was independently homogenised and a DNA extraction was performed using the DNeasy Blood and Tissue Spin-Column Kit (Qiagen). DNA extracts were quantified using Nanodrop 8000 UV–Vis Spectrophotometer (Thermo Scientific) and the corresponding subsample pairs were combined at a ratio of 1:10 in the amount of DNA for Coleoptera to Acari plus Collembola (according to the range of expected species diversity of these two fractions), in order to minimise the biomass bias in the sequencing depth of the two mesofauna components. For metabarcoding, the bc3’ fragment corresponding to 418 bp of the 3’ end of the COI barcode region was amplified. Primers included a tail corresponding to the Illumina P5 and P7 sequencing adapters for subsequent library preparation (see Arribas et al., 2016). For each sample, three independent PCR reactions were performed and the amplicons were pooled. All information regarding primers and PCR reagents and conditions is given in Table S2. Amplicon pools were cleaned using Ampure XP magnetic beads, and used as template for a limited-cycle secondary PCR amplification to add dual-index barcodes and the Illumina sequencing adapters (Nextera XT Index Kit; Illumina, San Diego, CA, USA). The resulting metabarcoding libraries were sequenced on an Illumina MiSeq sequencer (2 x 300 bp paired-end reads) on ~ 1% of the flow cell each, to produce paired reads (R1 and R2) with a given dual tag combination for each sample. Negative controls were maintained across all the different steps above and were sequenced as three independent metabarcoding libraries.

### Bioinformatics read processing

Raw reads were quality checked in Fastqc (Babraham Institute, 2013). Primers were trimmed using fastx_trimmer and reads were processed in Trimmomatic (Bolger, Lohse, & Usadel, 2014) using TRAILING:20. Based on results from (Andújar, Arribas, Gray, et al., 2018) on the test of multiple tools and parameters for diverse metazoan metabarcoding samples, we further processed each library independently following several steps of the Usearch (Edgar, 2013) pipeline: reads were merged (option mergepairs – −fastq_minovlen 50, −fastq_maxdiffs 15), quality-filtered (Maxee = 1), trimmed to full length amplicons of 418 bp (−sortbylength), dereplicated (−fastx_uniques) and denoised (−unoise3, −minsize 4). Denoised reads from the 48 libraries for each region, representing putative haplotypes, were combined and dereplicated to get a collection of unique sequences for each regional dataset. The surviving reads were assigned to high-level taxonomic categories with the lowest common ancestor (LCA) algorithm implemented in MEGAN V5 (Huson, Auch, Qi, & Schuster, 2007). Each read was subjected to BLAST searches (blastn-outfmt 5 - evalue 0.001) against a reference library including the NCBI *nt* database (Accessed December 2016) plus 382 sequences corresponding to Acari and Collembola collected at Sierra de Grazalema. BLAST matches were fed into MEGAN to compute the taxonomic affinity of each read. This high level taxonomic assignation allowed extracting reads corresponding to the three target groups Acari, Collembola and Coleoptera, while excluding other taxa present in the bulk samples. Reads corresponding to the target groups were then aligned in Geneious using MAFFT and the Translation Align option, and those with insertions, deletions or stop codons disrupting the reading frame were excluded.

Surviving haplotypes from each region were further filtered to remove likely nuclear mitochondrial (numts) pseudogenes, following a protocol based on the relative abundance of co-distributed reads (Andújar et al. under review). The set of putative haplotypes for Acari, Collembola and Coleoptera was used to generate a community table with read-counts (haplotype abundance) by sample against the complete collection of reads (i.e., reads before the dereplicating and denoising steps) using Usearch (− search_exact option). Using these abundances, we firstly removed from each library those haplotypes with four or fewer reads according to the criteria used for the denoising (see above). Next, we identified haplotypes that, in all the libraries where they were present, contributed less than 1% of the total reads of the library. All reads falling in this category were then removed from the analysis, as an auxiliary criterion to define spurious copies not representing the true mitochondrial haplotypes. The 1% cut-off value removes most of the spurious numts while maximizing the number of real haplotypes to be further analysed (Andújar et al. under review). Community tables of fully filtered haplotypes were then transformed into incidence (presence/absence) data, that added to the haplotype filtering before, resulted in normalised samples for further analyses.

### Analysis of community composition and assembly at multiple thresholds of genetic similarity

The set of filtered haplotypes was used to generate an UPGMA tree with corrected genetic distances (F84 model), and based on this tree all haplotypes were grouped into clusters of genetic similarity at different thresholds (1%, 2%, 3%, 4%, 5%, 6% and 8%). This grouping procedure based on patristic pairwise distances over a phylogenetic tree including all haplotype sequences provided multiple hierarchical levels that each can be used to estimate alpha diversity (Figure 1 provides a graphical abstract of the workflow). These diversity measures were estimated for the richness of lineages by sample for the whole mesofauna community but also for the subsets corresponding to Acari, Collembola and Coleoptera. To test for significant differences in alpha diversity between the communities of different habitats and soil layers of each sampling point, repeated-measures ANOVAs were conducted using habitat and soil layer as grouping factors and sampling point as a within-subjects factor. For each of the three local settings, total accumulative richness (local scale richness) by habitat and soil layer and the contribution of mites, springtails and beetles was also calculated for the various levels of genetic similarity. Endemicity by sampling point was computed for each hierarchical level (once DEEP and SUP samples were combined) as the lineages present exclusively at a single sampling point in the region divided by the total number of lineages found in the region. To assess whether the endemicity by sampling point differed between the communities of forests and grasslands, Wilcoxon tests were conducted using habitat as a grouping factor. For each of the three local settings, the local scale endemicity, defined as the lineages present exclusively at a particular sampling point divided by the total number of lineages in that community, was also calculated for the multiple levels of genetic similarity.

For the multi-hierarchical assessment of the variation in community composition at the local scale, the community dissimilarity matrices were generated for total beta diversity (Sorensen index, βsor) and its additive turnover (Simpson index, βsim) and nestedness (βsne) components (Baselga, 2010), for each level of genetic similarity. Community composition matrices were also used for non-parametric multidimensional scaling (NMDS) and plots were created with the *ordispider* option to visualise the compositional ordination of the communities according to the respective habitat and soil layer. To assess for significant differences, permutational ANOVAs were conducted over the community dissimilarity matrices using 999 permutations and the habitat and the soil layer as grouping factors and sampling point as a within-subjects factor. The significant relationships between the dissimilarity matrices generated for the Acari, Collembola and Coleoptera were assessed independently by permutational multiple regression on distance matrices (MRM), and additional NMDS ordinations and permutational ANOVAs as before were in this case conducted for each taxonomic group.

The analysis of the variation in community composition with spatial distance followed the ‘multi-hierarchical macroecology’ approach of (Baselga et al., 2013) which is based on the joint analysis of distance decay of similarity patterns across the different genetic levels. For each local setting and habitat, the relationship of community similarity between pairs of points (1 – pairwise beta diversity, see above) with their spatial distance (computed in kilometres as the Euclidean distance) was assessed independently at each level of genetic similarity (from haplotypes to 8% lineages). A negative exponential function was used to adjust a generalized linear model (GLM) with Simpson similarity as response variable, spatial distance as predictor, log link and Gaussian error, and maintaining the spatial distances untransformed (Gómez-Rodríguez & Baselga, 2018). Finally, the existence of a fractal pattern (power law function) in the distance-decay curves across the levels of genetic similarity was assessed by a log–log Pearson correlation of genetic level and, independently: (a) number of lineages, (b) initial similarity, and (c) mean similarity. High correlation values are indicative of self-similarity in lineage branching (i.e., number of lineages) and/or spatial geometry of lineage distributional ranges (i.e., initial and mean similarity, (Baselga et al., 2015), which are predicted under a neutral process of community evolution.

In an equivalent way, these analyses were also conducted to assess the relationships of community similarity and environmental distance as computed using Gower's distance over the recorded variables characterising the sampling points (Table S1). In the cases where this relationship was significant, variance partitioning was conducted to assess the fractions of variance in community dissimilarity that are uniquely and jointly explained by spatial and environmental distance. All analyses were performed using the R-packages vegan (Oksanen et al., 2013), cluster, PMCMR, hier.part, ecodist, and betapart (Baselga & Orme, 2012).

## RESULTS

### Multi-hierarchical assessment of alpha and gamma diversity of soil mesofauna

Processing of 144 soil samples using the FBF protocol, followed by double dual indexing of *cox1* amplicons and Illumina MiSeq sequencing, produced 51433 to 375211 sequence reads per sample for GRA, 11307 to 159244 for ALZ, and 43562 to 128149 for CUE. Filtering of raw reads using standard protocols of read curation and denoising, followed by removal of likely nuclear mitochondrial pseudogenes generated a conservative set of clean sequences representing the mitochondrial haplotypes. A total of 1124, 1009 and 992 haplotypes where found for the GRA, ALZ and CUE local areas respectively, and these numbers declined rapidly when haplotypes were grouped at increasing threshold values, e.g. 511, 479 and 480 lineages at 3% similarity (Table 1), but they declined only slightly further at the higher thresholds, indicating the point at which stable groups are obtained that broadly represent the species level. The relative proportions of mites, springtails and beetles were similar across the three local settings and hierarchical levels, with Acari representing the richest group (around the 50% of clusters) followed by Collembola and Coleoptera in similar proportions (Table 1).

**Table 1.**
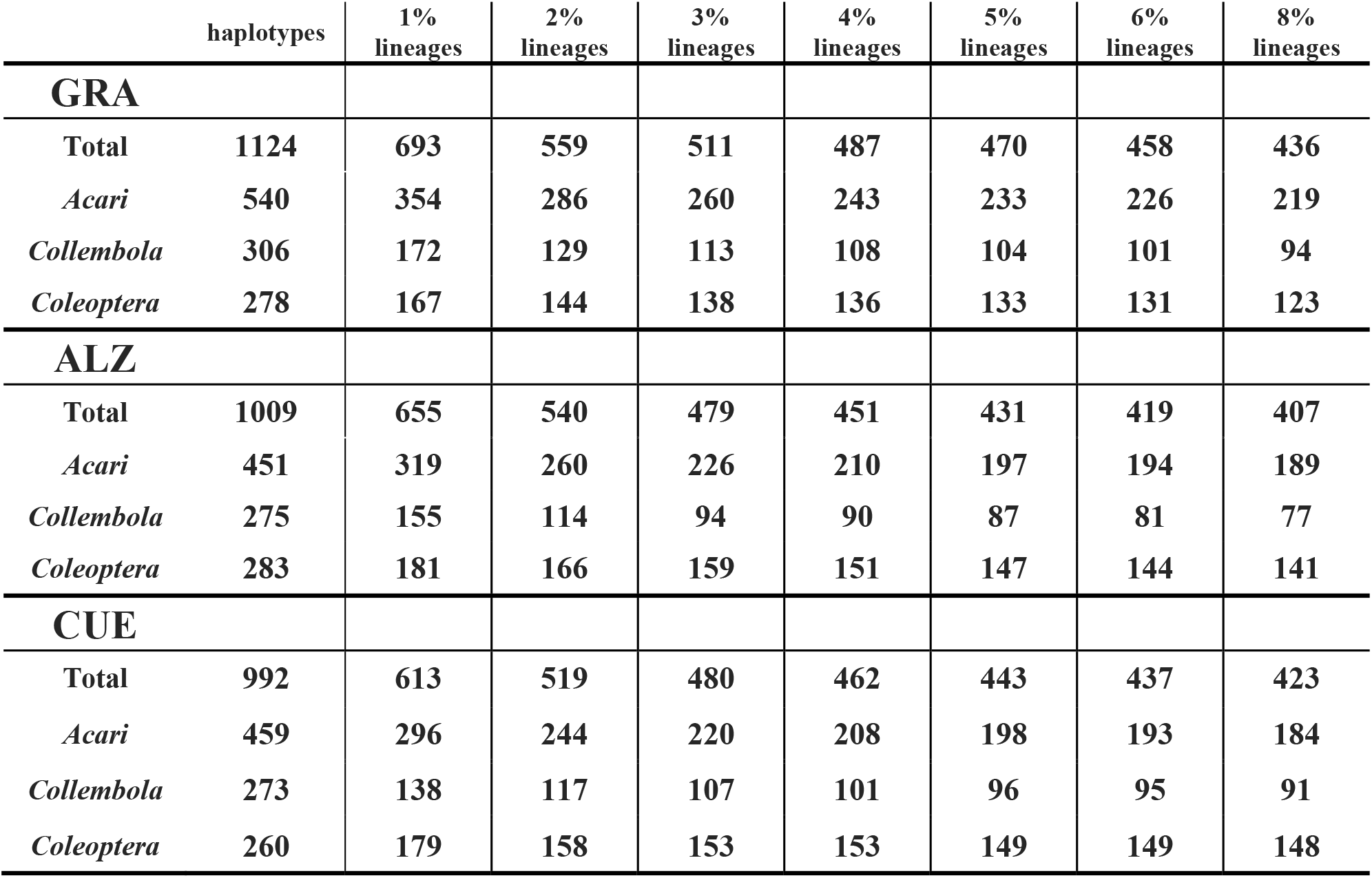
Total number of haplotypes and clusters for Acari, Collembola and Coleoptera for each local setting (GRA, ALZ, CUE) at increasing levels of genetic divergence thresholds.

The patterns of richness by sample (alpha diversity) for the different habitats, soil layers and genetic thresholds were similar for the three regions, with mean values between 35 - 60 haplotypes and 25 - 42 lineages at 3% per sample (Fig. 2 A, B, C and Fig. S1). Superficial soils had significantly higher diversity than their corresponding deep soil counterparts for the overall dataset and for both of the forest and grassland habitats assessed independently (Fig. 2 A, B, C, Fig S1 and Table S3). At GRA, forest habitat showed significantly higher alpha diversity per sample than grassland but no significant differences between forest and grassland were found for ALZ and CUE (Fig. 2 A, B, C and Fig S1, Table S3).

**Figure 2.**
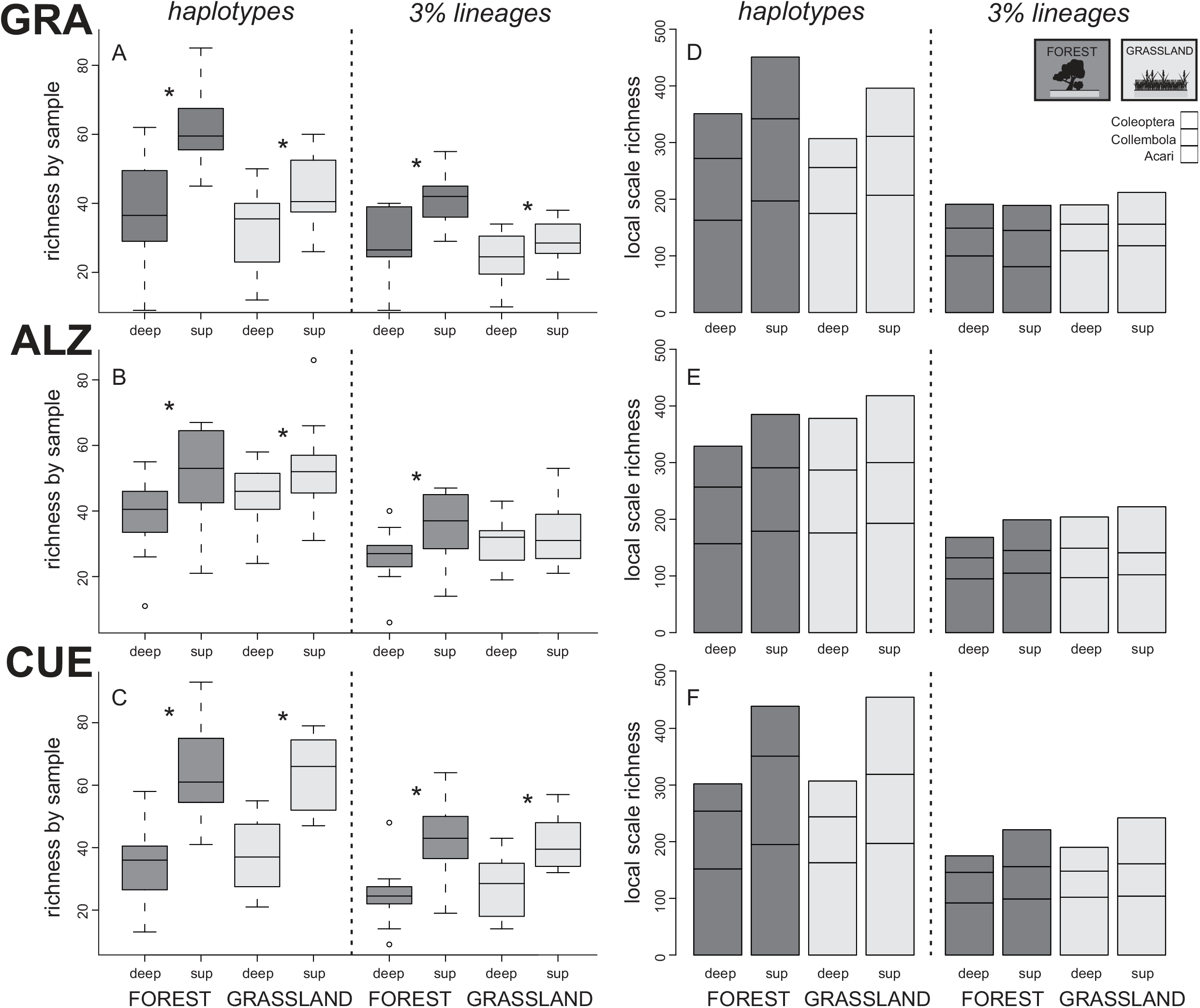
Richness of soil mesofauna lineages by sample (alpha diversity, A, B, C) and total accumulated richness (local scale richness, D, E, F) by habitat and soil layer for the three local settings (GRA, ALZ, CUE). Both measures are shown at the haplotype and the 3% genetic similarity levels. Forest habitat as dark grey, grassland habitat as pale gray, sup for superficial and deep for deep soil layers. Significantly different richness of lineages by sample (repeated-measures ANOVA *p* < 0.05) between deep and superficial communities of each habitat are indicated by asterisks within A, B, C panels. The contribution of Acari, Collembola and Coleoptera to the local scale richness are shown within D, E, F panels.

The local-scale cumulative richness at each region (gamma diversity) showed more diverse communities for the superficial compared with deep layers, and gamma diversity was generally higher for forest than grassland habitats, but the differences between both habitat types were lower than observed for alpha diversity. Thus, species accumulation was higher for the grassland than forest habitats, and the grassland superficial layers had the highest total richness of haplotypes for ALZ and CUE, and the highest for the three regions at the 3% similarity level (Fig. 2 D, E, F and Fig S2). Patterns of alpha and gamma diversity for the subsets of mites, springtails and beetles were similar (Fig. S2 and S3).

### Multi-hierarchical assessment of beta diversity and endemicity of soil mesofauna

Compositional dissimilarity of communities within each of the three regions was high and was dominated by lineage turnover *βsim*, instead of nestedness *βsne* (0.8 > *βsim* > 0.95), across all hierarchical levels. NMDS showed a consistent pattern of the forest and grassland habitats as the main driver of the ordination while soil layers had a secondary role (Fig. 3, Fig. S4). Accordingly, for the three regions and all genetic levels, the community composition was significantly different for both habitats and soil layers but the proportion of variance explained by the forest-grassland factor was always higher (Fig. 3 and Table S4). Beta diversity matrices for mites, springtails and beetles showed high and significant correlations for each of the genetic similarity levels (Table S5), and when independently analysed, these main taxonomic groups each showed similar patterns of community composition.

**Figure 3.**
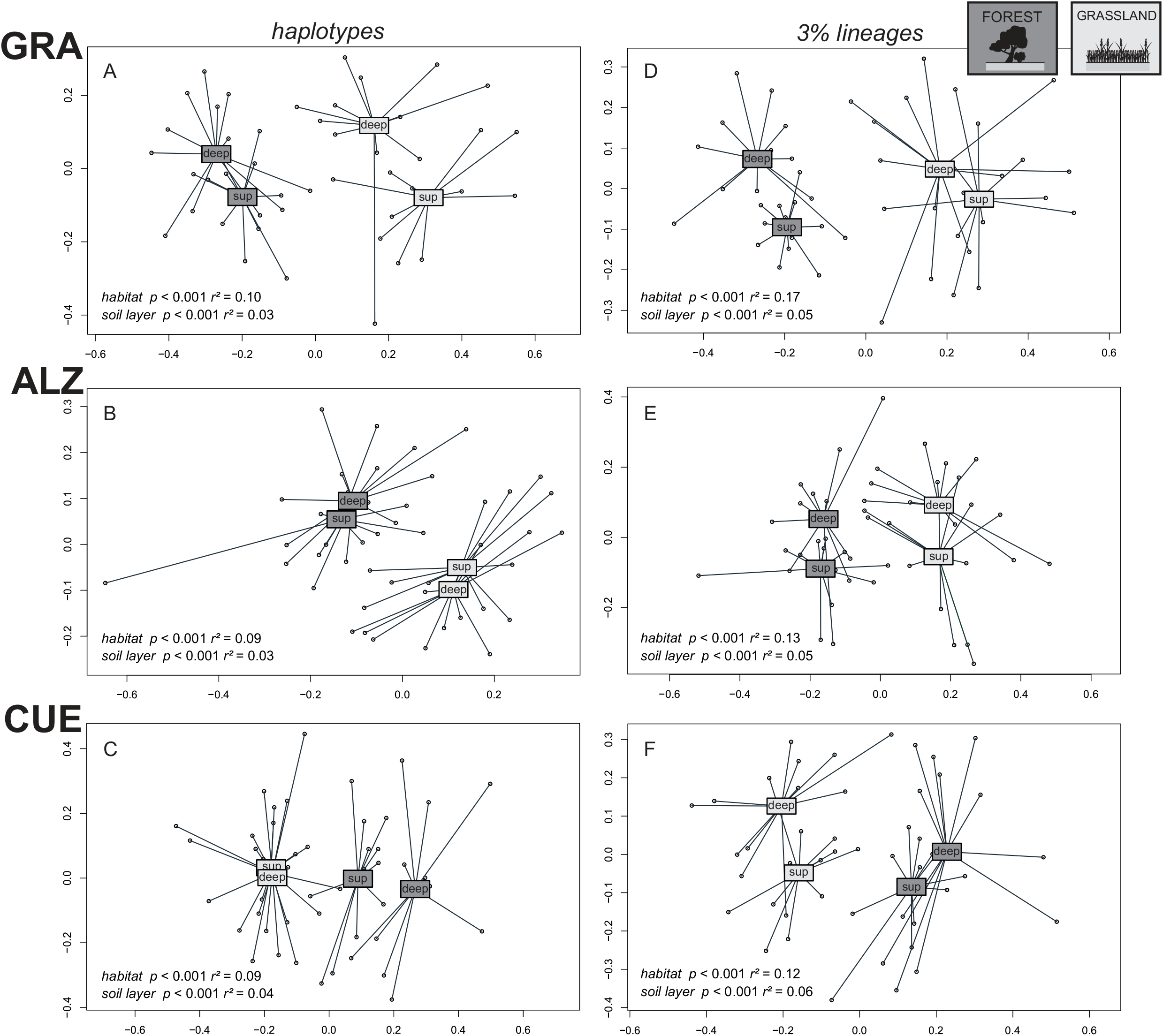
NMDS ordinations of the soil mesofauna samples according to the variation in community composition (Simpson index, βsim) within the three local settings (GRA, ALZ, CUE) and at the haplotype and the 3% genetic similarity levels. Forest habitat as dark grey, grassland habitat as pale grey, sup for superficial and deep for deep soil layers. Explained variation (*r^2^*) and significance (*p*) of each grouping factor from the permutational ANOVAs over the community dissimilarity matrixes are shown.

Community similarity (1-pairwise beta diversity) significantly decreased with spatial distance (distance decay) at all levels of genetic similarity for both the forest and grassland habitats, and these patterns were remarkably consistent across the three local settings (Fig. 4, Table S6). The slopes of the exponential decay curves were very similar at all threshold levels, and assemblage similarity increased with each level (Fig. 4, Table S6). The levels of genetic similarity showed a high and significant log–log correlation with the number of lineages (0.90 < *r^2^* > 0.96, *p* < 0.001), initial similarity (0.86 < *r^2^* > 0.96, *p* < 0.001) and mean similarity of communities (0.89 < *r^2^* > 0.95, *p* < 0.001) for all three regions and two habitats, as expected if community variation across genetic similarity levels can be described by a fractal geometry (Baselga et al., 2013, 2015).

**Figure 4.**
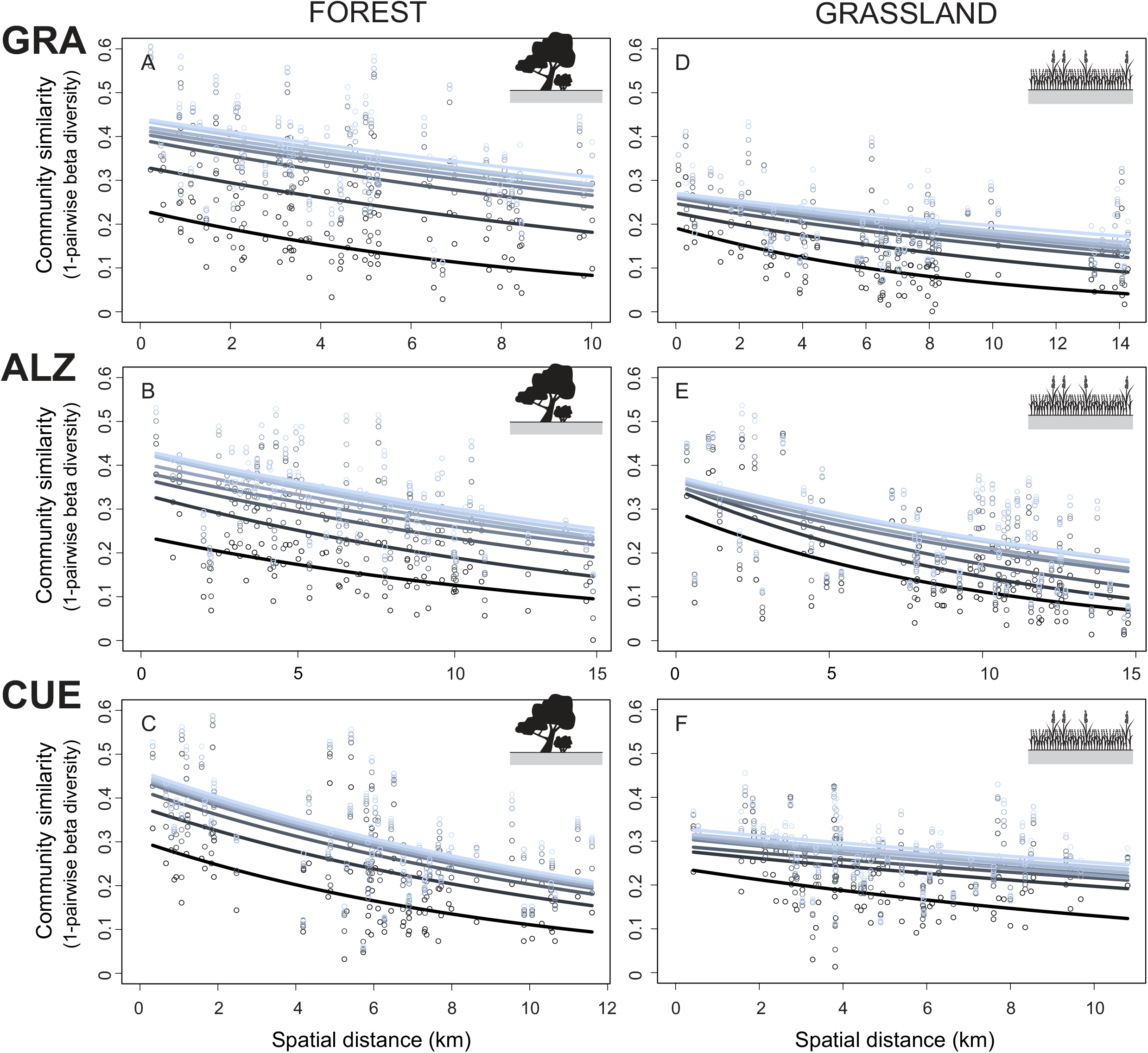
Distance decay of soil mesofauna community similarity at multiple levels of genetic similarity (from haplotype, black to 8% genetic similarity level, pale grey) within the three local settings (GRA, ALZ, CUE) and for forest and grassland habitats.

Comparisons of distance-decay relationships between forests and grasslands showed similar values for explained variance and for the slopes for the three regions (Fig. 4, Table S6). However, there was a consistent pattern of a lower initial community similarity in grasslands than in forests, particularly above the haplotype level (Fig. 4). Similarly, the local-scale (mean) dissimilarity of communities was always higher for grasslands than for forests and the differences between both increased across the levels of genetic similarity (Fig. 5 A, B, C). A decrease in community similarity with environmental distance was only significant in the case of the forests from ALZ and CUE, but variance partitioning showed that uniquely explained variance in environmental distance, i.e. independently of the spatial distance, was reduced (5 – 9 % of explained variation at all levels) compared with the uniquely explained variance in spatial distance (23 – 31 % of explained variation).

**Figure 5.**
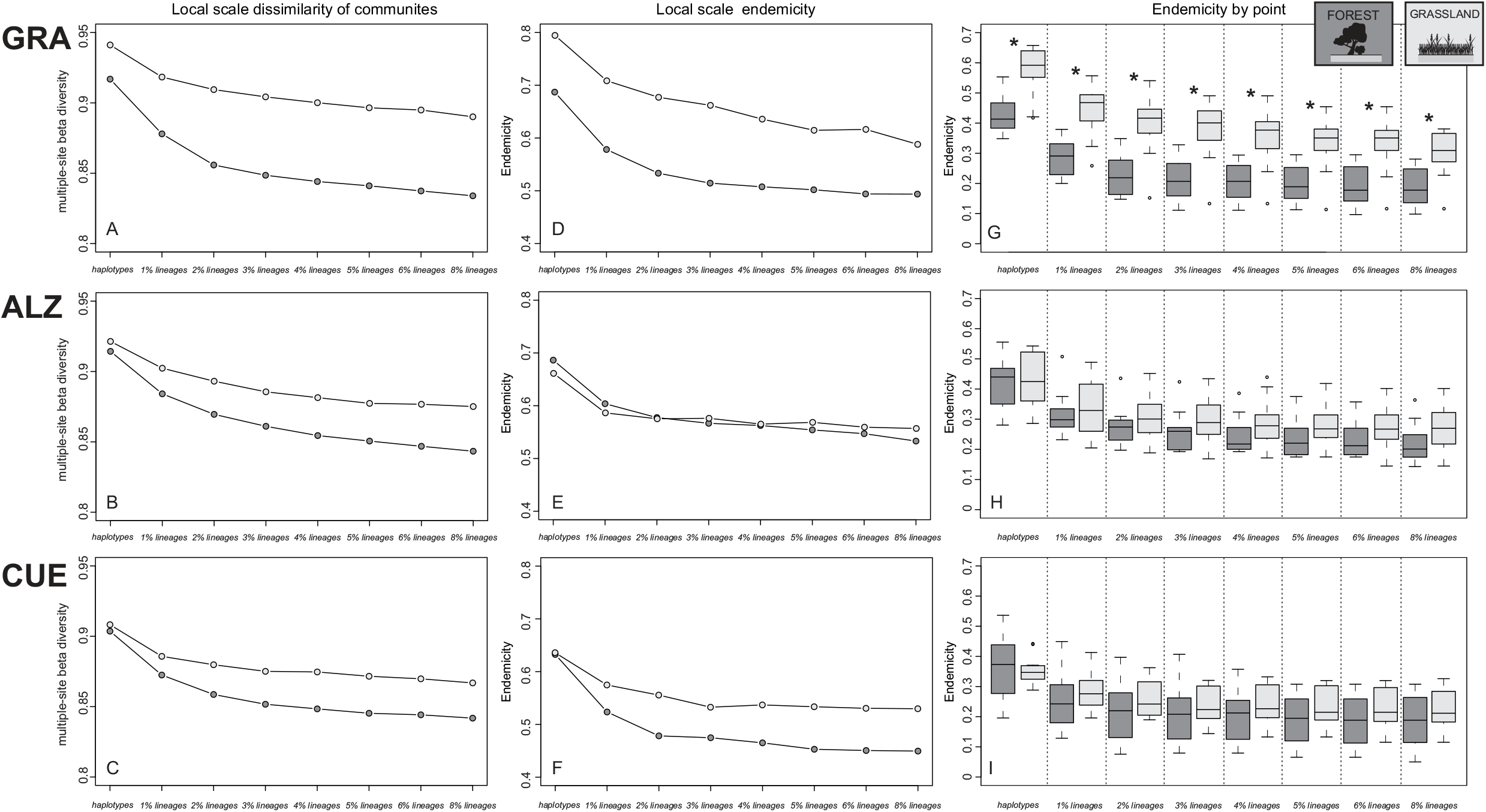
Dissimilarity of soil mesofauna communities (A, B, C), regional endemicity (lineages present exclusively at a single sampling point in the region divided by the total number of lineages found, D, E, F) and endemicity by sampling points (lineages present exclusively at a particular sampling point divided by the total number of lineages in that community, G, H, I) at multiple levels of genetic similarity within the three local settings (GRA, ALZ, CUE) and for forest (dark grey) and grassland (pale grey) habitats. Significantly different endemicity by sampling point (Wilcoxon tests *p* < 0.05) between forest and grassland communities at each hierarchical level is indicated by asterisks in panels G, H, I.

The endemicity within the GRA, ALZ and CUE regions ranged from 71%, 64% and 58% at the haplotype level to 55%, 53% and 46% for lineages at the 3% threshold, respectively (Table 1). Comparisons between forest and grassland habitats showed that the local scale endemicity of grassland communities was higher in the case of GRA and CUE but similar in both habitats for ALZ (Fig. 5 D, E, F). The endemicity by sampling points was consistently higher for grassland than for forest local communities particularly above the haplotype level, but the differences were significant only in the case of the GRA localities (Fig. 5 G, H, I).

## DISCUSSION

In total, soil samples from three Iberian mountain regions produced over 1000 species and nearly 3000 haplotypes. Their distribution was determined across numerous sampling points, demonstrating the power of mitochondrial cMBC to overcome previous impediments to studying the arthropod mesofauna of the soil using conventional morphological and molecular approaches. Data analysis in a multi-hierarchical framework revealed a strong spatial community structure and high levels of endemicity at haplotype, species and supra-specific levels, even at sampling points that were mostly within a few kilometres of each other (maximum 15 km). Patterns of turnover and endemicity were similar in all three independent study regions and in the grassland and forest biomes (that each harbour largely non-overlapping communities). Distance decay is evident at all hierarchical levels, and can be described as self-similar. The coincidence of community turnover at population level and species level is expected if soil arthropod assemblages are predominantly driven by distance-based parameters, if movement is strongly constrained at the local scale and over time. In addition, the overriding importance of habitat-related processes was apparent from the strong differentiation of grassland and forest communities, which again was seen recurrently in each of the three study regions.

The study extends existing comparative analyses of soil mesofauna by improving the taxonomic resolution, providing haplotype level variation and spanning most lineages composing the soil arthropod communities. Broad surveys of invertebrate soil diversity using HTS have commonly relied on markers of low species-level resolution and via eDNA extracted from small soil samples (Bahram et al., 2016; Wu et al., 2011; Zinger et al., 2019). Other studies have characterise specific groups of mites or springtails by individualised processing of specimens and relying on morphological assignment to generate species-level data (Caruso, Schaefer, Monson, & Keith, 2019; Ingimarsdóttir et al., 2012), but see also Young, Proctor, DeWaard, & Hebert (2019) on molecular species assignment. HTS data now greatly increase the potential of expanding both the number of species studied and the level of detail at which intra-specific variation for each is captured. To our knowledge, this is the first study that provides haplotype level data for entire communities (one square meter of leaf litter and humus and ca. 20 litres of soil per sampling point) of the three most species rich soil arthropods, which allows surveys of community composition and species turnover at an unprecedented level of detail, both spatially and genetically. With this data in hand, community level responses to distance-based parameters can be assessed that may be not evident in other types of studies. In addition, the combined haplotype and species level data permit the exploitation of the hierarchical framework of Baselga et al. (2013, 2015) for discriminating between distance- and niche-based factors of community assembly.

### The limited spatial scale of dispersal in soil arthropods

The literature exploring the community assemblage of arthropod mesofauna at the local scale is generally arguing for the selection by abiotic and/or biotic environmental factors as the predominant mechanisms (see Berg, 2012; Thakur et al., 2019 for a recent review) but stochastic and purely spatial patterns have also been reported, pointing to a contribution of dispersal and demographic processes at least in some local settings (Caruso et al., 2012; Gao et al., 2014; Gao, Liu, Lin, & Wu, 2016; Zinger et al., 2019). However, strong dispersal constraints have rarely been recognised as an important mechanism of soil mesofauna assembly at the local scale, and this could be in part due to the potentially low taxonomic resolution of most of the community-level studies in this group that used morphological or 18S rRNA gene species assignment (see Tang et al., 2012 on the low taxonomic resolution of this marker). Our results demonstrate high community differentiation at the kilometre scale for both genetic and species levels. The key observation from the multi-hierarchical analysis is the correlated distance decay at haplotype and species level. Self-similarity is expected to be eroded by selection on adaptive traits at the species level, but not at the (neutral) haplotype level (Baselga et al., 2015; Gómez-Rodríguez et al., 2018). As the data largely confirm the self-similarity of distance decay at haplotype and species level, this is interpreted to support the predominant role of dispersal limitation driving community assembly. The predominance of the dispersal constraints seems to emerge at short spatial distances within the soil matrix, and the evident high turnover with physical distance suggests that our sampling within each study regions (local scale) is beyond the scale of a single metacommunity. Short dispersal distances probably have affected a significant proportion of lineages within these communities over evolutionary timescales in a largely stable spatial setting. The spatiotemporal continuum expected under this scenario predicts that lineages in more distant places have diverged at a more distant time point in evolutionary history (Baselga et al., 2013, 2015), and our findings of a largely regular distance decay at higher levels are consistent with this prediction. Additional evidence for the role of short dispersal distance driving the local community assembly comes from the high microendemicity found at all hierarchical levels, an overall picture which is not expected under a scenario with predominant environmental drivers nor ecological drift without dispersal limitation.

Yet, the influence of environmental drivers cannot be discarded entirely. Even if the recorded environmental variables did not explain the variation in community composition, a significant portion of the unexplained variance in the distance-decay curves potentially suggests the influence of non-spatial factors determining the community composition. Edaphic parameters such as soil pH or organic matter have been shown to explain a significant part of the variance observed in the distribution of the soil mesofauna communities (Caruso et al., 2012; Gao et al., 2014), and here could be driving at least part of the unexplained variation within the different habitats and regions. Edaphic environmental variables are often spatially structured and so have been also reported as potential drivers of purely spatial patterns in mesofauna communities (Caruso et al., 2019, 2012). However as exposed before, this possibility is poorly supported here, as similar spatial structures were independently found within the different habitats and regions, mirroring the distance-decay patterns at the (neutral) haplotype level, and hence suggesting that dispersal limitation is the main driver of the local spatial structure of the studied mesofauna communities.

The small spatial scale of turnover and endemicity is consistent with population genetic studies in soil mesofauna showing deep genetic breaks even over relative short geographic distances (Andújar, Pérez-González, et al., 2017; Cicconardi et al., 2013). In contrast, our results are not concordant with the extended view of long-distance dispersal as prevalent process for soil mesofauna, as might be expected from evidences of passive dispersal by air, water or in marine plankton (Decaëns, 2010; Thakur et al., 2019; Wardle, 2002). Existing reports of long-distance dispersal are mainly into virgin isolated habitats (Ingimarsdóttir et al., 2012) or recently deglaciated areas (Gan et al., 2019), or may involve the detection of mesofauna during transport (Coulson, Hodkinson, & Webb, 2003; Schuppenhauer et al., 2019). However, they do not inform about colonisation and establishment success (effective dispersal) and possibly only pertain to a few highly dispersive species. Additionally, the dispersal potential may have been overestimated due to the low resolution of morphological species identification (Cicconardi et al., 2013) leading to perceived low turnover among sites, as evident from recent large-scale barcoding studies (Young et al., 2019). Our results at the community level thus raise doubts about a generalised dispersal advantage for small-bodied arthropods and instead indicate very small dispersal distances, even over evolutionary time scales, for the majority of species that make up the complex mesofauna communities of the soil. This scale and dynamics of community assembly contrasts with patterns and processes reported for the microbial eukaryote diversity of the soil (Bahram et al., 2016) and aligns with recent empirical evidences suggesting that at the local scale dispersal rates may be much lower for soil mesofauna than for microfauna (Zinger et al., 2019). In the context of the overall arthropod diversity (for which soil mesofauna comprises the majority of the smallest fraction), our results are not supporting the macroecological prediction for a reduce impact of dispersal limitation in the assemblage for small-bodied components compared with their bigger counterparts (de Bie et al., 2012; Ricklefs, 2004) and highlight the uniqueness of ecological and evolutionary processes driving the biodiversity of these edaphic arthropods (Andújar, Arribas, & Vogler, 2017; Andújar, Pérez-González, et al., 2017).

### The role of dispersal constraints within a habitat-based framework

In spite of the evidence for an important role for dispersal limitation within each habitat type, the greatest effect was the differentiation of grassland and forest communities, which share very few species even in close (meters) spatial proximity. Previous studies also have shown great differences in soil arthropod community composition between beech forest and adjacent grassland (Caruso et al., 2012), and twice higher species richness in the forest community. The grassland-forest dichotomy in community composition is concordant with these findings, but the total diversity in either type of community was more complex: alpha diversity tended to be higher for forest habitats, but lineage accumulation across multiple sites was higher for the grasslands, resulting in higher overall landscape richness (gamma diversity). Grassland communities also had consistently lower initial and mean community similarities in the corresponding distance decay curves, together with higher levels of both point and local scale endemicity. These results are recurrent across the three sampling areas and point to slightly higher long-term dispersal constraints for the mesofauna in the grasslands studied.

Grassland species are expected to experience higher environmental variability and greater extremes, which are moderated within forested patches and thus are presumably more stable (De Frenne et al., 2019). Under the habitat stability hypothesis (Ribera & Vogler, 2000; Southwood, 1977), low species turnover is predicted in less stable habitats due to the stronger positive selection on traits promoting dispersal that are required to persist in ephemeral environments. However, our findings are not aligned with this hypothesis, suggesting similar local-scale patterns of lineage turnover within both habitats and with slightly stronger community structure for the presumably less stable grasslands. Further studies comparing the assemblages of both habitat types across gradients of stability (e.g. latitude) are fundamental to identify the similarities and disparities in the processes driving the diversity and structure of the mesofauna communities. Regardless of the potential differences between the two habitat types, the high overall turnover suggests the long-term stability of the soil environment without which the high spatial structure at multiple hierarchical levels and between grassland and forest habitats could not have arisen.

### Extrapolating beyond the local scale

Disentangling the mechanisms at larger spatial scales might be challenging without the understanding of the underlying processes at relatively fine scales. The recurrence of the local patterns in each of the three study regions and across the three major taxonomic groups, corroborates the hypothesis of an underlying process of stochastic dispersal of individuals, affected by a universal type of dispersal constraint. This seems to affect to a majority of the soil arthropod fauna composing these communities, regardless of their taxonomic affinity, species traits or functional role. The soil matrix provides a common sphere in which these processes are played out, and if these soils are similar, complex communities, on average, appear to respond in a similar way. The two habitat types clearly provide a different overall setting, obvious from the very different species present, but they also impact the respective species pool in similar ways. With the local-scale patterns and likely underlying processes reported here, questions arise about the impact for cross-regional and global spatial scales, and how these patterns and processes compare to aboveground arthropod components. If generalized across broader geographical scales and latitudes, the proposed process of a very reduced spatial scale of dispersal in soil mesofauna communities could be a major contribution to the overall gamma diversity of ecosystems and would suppose a great underestimation of global diversity on Earth. In this sense, further developments on the multi-hierarchical analysis of genetic and higher-level diversity from metabarcoding data has the potential to propel the characterisation of edaphic macrobial community structure into a new era of biodiversity discovery. By taking advantage of the full breadth of contemporary metabarcoding data at unprecedented taxonomic and geographic scales, the advances made here will be able to provide unique insights into the ecological and evolutionary processes that determine the magnitude and spatial distribution of soil arthropods.

## ACKNOWLEDGMENTS

This research was funded by Newton International Program, UK to PA and the NHM Biodiversity Initiative to APV. PA was supported by two postdoctoral grants from the Royal Society (Newton International Program, UK) and the Spanish Ministry of Economy and Competitiveness (MINECO, Spain) within the Juan de la Cierva Formación Program. CA was supported by the Spanish Ministry of Economy and Competitiveness (MINECO, Spain) (CGL2015-74178-JIN). Thanks are due to the regional government of Andalucía and Castilla-la-Mancha (Spain); Alex Aitken, Andie Hall, Stephen Russell and Peter Foster (all NHM) for their technical assistance and to Jesús Arribas who assisted with field sampling.

## AUTHOR CONTRIBUTIONS

Statement of authorship: P.A., C.A. and A.P.V. conceived the work; P.A. and C.A. collected and analysed the data; P.A led the writing and all authors contributed to the discussion of results and the writing.

