## Supporting Information for "The limited spatial scale of dispersal in soil arthropods revealed with whole-community haplotype-level metabarcoding"

**Table S1.** Sampling point information associated to each soil mesofauna community sample.

| *Sample_code* | *Region* | *Soil_layer* | *Habitat* | *Latitude* | *Longitude* | *Elevation* | *Slope* | *Orientation* | *Stream prox* | *Site* | *Stones_sup* | *Vegetation* | *Humus* | *Stones_deep* | *Sand/Clay* | *Porosity* | *Roots* | *Temperature* | *Humidity* |
|---|---|---|---|---|---|---|---|---|---|---|---|---|---|---|---|---|---|---|---|
| ALZ_S1_D_F_A1 | ALZ | DEEP | Forest | 39.079052 | -1.316559 | 1065 | moderate | e | no | tree | many | yes | 10 | many | clay | normal | normal | 11 | 17 |
| ALZ_S1_S_F_A4 | ALZ | SUP | Forest | 39.079052 | -1.316559 | 1065 | moderate | e | no | tree | many | yes | 10 | many | clay | normal | normal | 11 | 17 |
| ALZ_S10_D_G_B2 | ALZ | DEEP | Grassland | 39.069801 | -1.363245 | 943 | moderate | e | yes | free_soil | no | yes | 0 | few | sand | high | many | 12 | 8.5 |
| ALZ_S10_S_G_B5 | ALZ | SUP | Grassland | 39.069801 | -1.363245 | 943 | moderate | e | yes | free_soil | no | yes | 0 | few | sand | high | many | 12 | 8.5 |
| ALZ_S11_D_F_C2 | ALZ | DEEP | Forest | 39.061705 | -1.364947 | 949 | no | e | no | free_soil | few | yes | 5 | many | sand | normal | many | 11 | 10.5 |
| ALZ_S11_S_F_C5 | ALZ | SUP | Forest | 39.061705 | -1.364947 | 949 | no | e | no | free_soil | few | yes | 5 | many | sand | normal | many | 11 | 10.5 |
| ALZ_S12_D_F_D2 | ALZ | DEEP | Forest | 39.041376 | -1.342276 | 996 | no | s | no | tree | few | yes | 10 | no | sand | high | normal | 11 | 11.5 |
| ALZ_S12_S_F_D5 | ALZ | SUP | Forest | 39.041376 | -1.342276 | 996 | no | s | no | tree | few | yes | 10 | no | sand | high | normal | 11 | 11.5 |
| ALZ_S13_D_F_E2 | ALZ | DEEP | Forest | 39.03714 | -1.341109 | 987 | no | se | no | tree | no | yes | 15 | no | sand | high | many | 10 | 9.5 |
| ALZ_S13_S_F_E5 | ALZ | SUP | Forest | 39.03714 | -1.341109 | 987 | no | se | no | tree | no | yes | 15 | no | sand | high | many | 10 | 9.5 |
| ALZ_S14_D_G_F2 | ALZ | DEEP | Grassland | 39.015237 | -1.257104 | 1012 | high | sw | yes | batter | no | yes | 0 | no | sand | normal | normal | 12 | 15 |
| ALZ_S14_S_G_F5 | ALZ | SUP | Grassland | 39.015237 | -1.257104 | 1012 | high | sw | yes | batter | no | yes | 0 | no | sand | normal | normal | 12 | 15 |
| ALZ_S15_D_F_G2 | ALZ | DEEP | Forest | 39.073591 | -1.326018 | 1050 | moderate | e | no | tree | few | yes | 10 | many | clay | normal | many | 11 | 22 |
| ALZ_S15_S_F_G5 | ALZ | SUP | Forest | 39.073591 | -1.326018 | 1050 | moderate | e | no | tree | few | yes | 10 | many | clay | normal | many | 11 | 22 |
| ALZ_S16_D_F_H2 | ALZ | DEEP | Forest | 39.116978 | -1.268721 | 890 | no | n | no | tree | few | yes | 10 | no | sand | high | many | 12 | 7.5 |
| ALZ_S16_S_F_H5 | ALZ | SUP | Forest | 39.116978 | -1.268721 | 890 | no | n | no | tree | few | yes | 10 | no | sand | high | many | 12 | 7.5 |
| ALZ_S17_D_G_A3 | ALZ | DEEP | Grassland | 39.077077 | -1.341129 | 940 | high | w | no | free_soil | no | yes | 0 | many | sand | high | many | 12 | 14 |
| ALZ_S17_S_G_A6 | ALZ | SUP | Grassland | 39.077077 | -1.341129 | 940 | high | w | no | free_soil | no | yes | 0 | many | sand | high | many | 12 | 14 |
| ALZ_S18_D_F_B3 | ALZ | DEEP | Forest | 39.014641 | -1.32426 | 932 | moderate | n | no | free_soil | few | yes | 5 | many | sand | normal | many | 11 | 7.5 |
| ALZ_S18_S_F_B6 | ALZ | SUP | Forest | 39.014641 | -1.32426 | 932 | moderate | n | no | free_soil | few | yes | 5 | many | sand | normal | many | 11 | 7.5 |
| ALZ_S19_D_G_C3 | ALZ | DEEP | Grassland | 39.009839 | -1.313442 | 923 | no | sw | no | batter | no | yes | 0 | many | sand | normal | many | 16 | 7 |
| ALZ_S19_S_G_C6 | ALZ | SUP | Grassland | 39.009839 | -1.313442 | 923 | no | sw | no | batter | no | yes | 0 | many | sand | normal | many | 16 | 7 |
| ALZ_S2_D_F_B1 | ALZ | DEEP | Forest | 39.079003 | -1.269371 | 1109 | no | se | no | tree | few | yes | 10 | many | clay | normal | many | 10 | 19.5 |
| ALZ_S2_S_F_B4 | ALZ | SUP | Forest | 39.079003 | -1.269371 | 1109 | no | se | no | tree | few | yes | 10 | many | clay | normal | many | 10 | 19.5 |
| ALZ_S20_D_G_D3 | ALZ | DEEP | Grassland | 38.991533 | -1.294382 | 888 | no | s | yes | batter | no | yes | 0 | few | sand | normal | many | 13 | 21 |
| ALZ_S20_S_G_D6 | ALZ | SUP | Grassland | 38.991533 | -1.294382 | 888 | no | s | yes | batter | no | yes | 0 | few | sand | normal | many | 13 | 21 |
| ALZ_S21_D_F_E3 | ALZ | DEEP | Forest | 38.990736 | -1.294465 | 894 | high | e | yes | free_soil | no | yes | 3 | few | sand | high | few | 12 | 8 |
| ALZ_S21_S_F_E6 | ALZ | SUP | Forest | 38.990736 | -1.294465 | 894 | high | e | yes | free_soil | no | yes | 3 | few | sand | high | few | 12 | 8 |
| ALZ_S22_D_F_F3 | ALZ | DEEP | Forest | 39.005323 | -1.353562 | 1008 | moderate | w | no | free_soil | no | yes | 5 | many | sand | low | normal | 11.5 | 10 |
| ALZ_S22_S_F_F6 | ALZ | SUP | Forest | 39.005323 | -1.353562 | 1008 | moderate | w | no | free_soil | no | yes | 5 | many | sand | low | normal | 11.5 | 10 |

| | | | | | | | | | | | | | | | | | | | |
|---|---|---|---|---|---|---|---|---|---|---|---|---|---|---|---|---|---|---|---|
| ALZ_S23_D_G_G3 | ALZ | DEEP | Grassland | 39.006774 | -1.353517 | 1000 | moderate | w | no | free_soil | no | yes | 0 | few | sand | normal | normal | 16 | 11 |
| ALZ_S23_S_G_G6 | ALZ | SUP | Grassland | 39.006774 | -1.353517 | 1000 | moderate | w | no | free_soil | no | yes | 0 | few | sand | normal | normal | 16 | 11 |
| ALZ_S24_D_F_H3 | ALZ | DEEP | Forest | 38.997231 | -1.329939 | 994 | no | ne | no | tree | few | yes | 7 | few | sand | normal | few | 13 | 10 |
| ALZ_S24_S_F_H6 | ALZ | SUP | Forest | 38.997231 | -1.329939 | 994 | no | ne | no | tree | few | yes | 7 | few | sand | normal | few | 13 | 10 |
| ALZ_S3_D_G_C1 | ALZ | DEEP | Grassland | 39.087363 | -1.21813 | 983 | moderate | n | no | batter | no | yes | 0 | no | clay | high | many | 11 | 10 |
| ALZ_S3_S_G_C4 | ALZ | SUP | Grassland | 39.087363 | -1.21813 | 983 | moderate | n | no | batter | no | yes | 0 | no | clay | high | many | 11 | 10 |
| ALZ_S4_D_G_D1 | ALZ | DEEP | Grassland | 39.088979 | -1.221653 | 977 | moderate | n | no | free_soil | no | yes | 0 | few | sand | high | many | 12 | 12 |
| ALZ_S4_S_G_D4 | ALZ | SUP | Grassland | 39.088979 | -1.221653 | 977 | moderate | n | no | free_soil | no | yes | 0 | few | sand | high | many | 12 | 12 |
| ALZ_S5_D_G_E1 | ALZ | DEEP | Grassland | 39.082192 | -1.230339 | 1088 | moderate | w | yes | free_soil | no | yes | 0 | many | clay | normal | many | 11 | 15 |
| ALZ_S5_S_G_E4 | ALZ | SUP | Grassland | 39.082192 | -1.230339 | 1088 | moderate | w | yes | free_soil | no | yes | 0 | many | clay | normal | many | 11 | 15 |
| ALZ_S6_D_G_F1 | ALZ | DEEP | Grassland | 39.112816 | -1.262565 | 894 | high | w | yes | batter | no | yes | 0 | no | sand | high | many | 12 | 9 |
| ALZ_S6_S_G_F4 | ALZ | SUP | Grassland | 39.112816 | -1.262565 | 894 | high | w | yes | batter | no | yes | 0 | no | sand | high | many | 12 | 9 |
| ALZ_S7_D_G_G1 | ALZ | DEEP | Grassland | 39.089842 | -1.343214 | 918 | moderate | w | yes | free_soil | few | yes | 0 | many | sand | normal | many | 12 | 15.5 |
| ALZ_S7_S_G_G4 | ALZ | SUP | Grassland | 39.089842 | -1.343214 | 918 | moderate | w | yes | free_soil | few | yes | 0 | many | sand | normal | many | 12 | 15.5 |
| ALZ_S8_D_F_H1 | ALZ | DEEP | Forest | 38.998332 | -1.304706 | 917 | moderate | e | no | free_soil | few | yes | 5 | few | sand | normal | normal | 12 | 7.5 |
| ALZ_S8_S_F_H4 | ALZ | SUP | Forest | 38.998332 | -1.304706 | 917 | moderate | e | no | free_soil | few | yes | 5 | few | sand | normal | normal | 12 | 7.5 |
| ALZ_S9_D_G_A2 | ALZ | DEEP | Grassland | 39.074887 | -1.365851 | 915 | no | sw | no | free_soil | few | yes | 0 | few | sand | normal | many | 12 | 18 |
| ALZ_S9_S_G_A5 | ALZ | SUP | Grassland | 39.074887 | -1.365851 | 915 | no | sw | no | free_soil | few | yes | 0 | few | sand | normal | many | 12 | 18 |
| CUE_S1_D_G_A7 | CUE | DEEP | Grassland | 40.469907 | -2.450134 | 773 | no | ne | yes | free_soil | no | yes | 0 | no | sand | normal | many | 13 | 19 |
| CUE_S1_S_G_A10 | CUE | SUP | Grassland | 40.469907 | -2.450134 | 773 | no | ne | yes | free_soil | no | yes | 0 | no | sand | normal | many | 13 | 19 |
| CUE_S10_D_F_B8 | CUE | DEEP | Forest | 40.523473 | -2.44588 | 1063 | moderate | s | no | tree | no | yes | 10 | many | sand | normal | normal | 11.5 | 15 |
| CUE_S10_S_F_B11 | CUE | SUP | Forest | 40.523473 | -2.44588 | 1063 | moderate | s | no | tree | no | yes | 10 | many | sand | normal | normal | 11.5 | 15 |
| CUE_S11_D_G_C8 | CUE | DEEP | Grassland | 40.534044 | -2.460185 | 894 | moderate | n | no | free_soil | no | yes | 0 | no | sand | normal | normal | 12 | 23 |
| CUE_S11_S_G_C11 | CUE | SUP | Grassland | 40.534044 | -2.460185 | 894 | moderate | n | no | free_soil | no | yes | 0 | no | sand | normal | normal | 12 | 23 |
| CUE_S12_D_F_D8 | CUE | DEEP | Forest | 40.532273 | -2.440537 | 1017 | high | n | no | batter | few | yes | 5 | many | clay | normal | normal | 11 | 24.5 |
| CUE_S12_S_F_D11 | CUE | SUP | Forest | 40.532273 | -2.440537 | 1017 | high | n | no | batter | few | yes | 5 | many | clay | normal | normal | 11 | 24.5 |
| CUE_S13_D_F_E8 | CUE | DEEP | Forest | 40.535201 | -2.433642 | 1135 | no | sw | no | free_soil | many | yes | 10 | many | clay | normal | normal | 12 | 16 |
| CUE_S13_S_F_E11 | CUE | SUP | Forest | 40.535201 | -2.433642 | 1135 | no | sw | no | free_soil | many | yes | 10 | many | clay | normal | normal | 12 | 16 |
| CUE_S14_D_F_F8 | CUE | DEEP | Forest | 40.571493 | -2.41474 | 1131 | no | n | no | tree | no | yes | 5 | few | clay | normal | normal | 11 | 15 |
| CUE_S14_S_F_F11 | CUE | SUP | Forest | 40.571493 | -2.41474 | 1131 | no | n | no | tree | no | yes | 5 | few | clay | normal | normal | 11 | 15 |
| CUE_S15_D_G_G8 | CUE | DEEP | Grassland | 40.563963 | -2.417689 | 1123 | moderate | w | no | free_soil | no | yes | 0 | many | clay | normal | normal | 12 | 23 |
| CUE_S15_S_G_G11 | CUE | SUP | Grassland | 40.563963 | -2.417689 | 1123 | moderate | w | no | free_soil | no | yes | 0 | many | clay | normal | normal | 12 | 23 |
| CUE_S16_D_G_H8 | CUE | DEEP | Grassland | 40.55306 | -2.434536 | 913 | no | n | no | free_soil | no | yes | 0 | no | sand | normal | many | 12 | 19 |

| | | | | | | | | | | | | | | | | | | |
|---|---|---|---|---|---|---|---|---|---|---|---|---|---|---|---|---|---|---|
| CUE_S16_S_G_H11 | CUE | SUP | Grassland | 40.55306 | -2.434536 | 913 | no | n | no | free_soil | no | yes | 0 | no | sand | normal | many | 12 | 19 |
| CUE_S17_D_F_A9 | CUE | DEEP | Forest | 40.484993 | -2.370474 | 754 | high | ne | yes | batter | few | yes | 5 | few | sand | normal | many | 13 | 15 |
| CUE_S17_S_F_A12 | CUE | SUP | Forest | 40.484993 | -2.370474 | 754 | high | ne | yes | batter | few | yes | 5 | few | sand | normal | many | 13 | 15 |
| CUE_S18_D_G_B9 | CUE | DEEP | Grassland | 40.511072 | -2.435947 | 930 | no | s | no | free_soil | no | yes | 0 | no | sand | normal | normal | 11 | 19 |
| CUE_S18_S_G_B12 | CUE | SUP | Grassland | 40.511072 | -2.435947 | 930 | no | s | no | free_soil | no | yes | 0 | no | sand | normal | normal | 11 | 19 |
| CUE_S19_D_G_C9 | CUE | DEEP | Grassland | 40.537468 | -2.464701 | 862 | no | s | no | free_soil | no | yes | 0 | no | sand | normal | many | 13 | 22 |
| CUE_S19_S_G_C12 | CUE | SUP | Grassland | 40.537468 | -2.464701 | 862 | no | s | no | free_soil | no | yes | 0 | no | sand | normal | many | 13 | 22 |
| CUE_S2_D_F_B7 | CUE | DEEP | Forest | 40.47045 | -2.449459 | 775 | moderate | w | no | tree | no | yes | 5 | few | sand | high | many | 13 | 5 |
| CUE_S2_S_F_B10 | CUE | SUP | Forest | 40.47045 | -2.449459 | 775 | moderate | w | no | tree | no | yes | 5 | few | sand | high | many | 13 | 5 |
| CUE_S20_D_F_D9 | CUE | DEEP | Forest | 40.534193 | -2.462847 | 878 | high | n | no | batter | no | yes | 5 | few | sand | normal | many | 11 | 17 |
| CUE_S20_S_F_D12 | CUE | SUP | Forest | 40.534193 | -2.462847 | 878 | high | n | no | batter | no | yes | 5 | few | sand | normal | many | 11 | 17 |
| CUE_S21_D_G_E9 | CUE | DEEP | Grassland | 40.529489 | -2.421068 | 947 | moderate | w | yes | batter | no | yes | 0 | no | clay | normal | normal | 13 | 18.5 |
| CUE_S21_S_G_E12 | CUE | SUP | Grassland | 40.529489 | -2.421068 | 947 | moderate | w | yes | batter | no | yes | 0 | no | clay | normal | normal | 13 | 18.5 |
| CUE_S22_D_G_F9 | CUE | DEEP | Grassland | 40.539091 | -2.407464 | 896 | moderate | s | yes | batter | few | yes | 0 | few | sand | normal | many | 15 | 18 |
| CUE_S22_S_G_F12 | CUE | SUP | Grassland | 40.539091 | -2.407464 | 896 | moderate | s | yes | batter | few | yes | 0 | few | sand | normal | many | 15 | 18 |
| CUE_S23_D_G_G9 | CUE | DEEP | Grassland | 40.53187 | -2.390288 | 810 | moderate | s | yes | free_soil | no | yes | 0 | no | sand | normal | many | 12 | 16.5 |
| CUE_S23_S_G_G12 | CUE | SUP | Grassland | 40.53187 | -2.390288 | 810 | moderate | s | yes | free_soil | no | yes | 0 | no | sand | normal | many | 12 | 16.5 |
| CUE_S24_D_F_H9 | CUE | DEEP | Forest | 40.537252 | -2.360959 | 845 | no | ne | no | tree | no | yes | 3 | few | sand | normal | normal | 13 | 30 |
| CUE_S24_S_F_H12 | CUE | SUP | Forest | 40.537252 | -2.360959 | 845 | no | ne | no | tree | no | yes | 3 | few | sand | normal | normal | 13 | 30 |
| CUE_S3_D_F_C7 | CUE | DEEP | Forest | 40.475659 | -2.457285 | 793 | moderate | ne | no | tree | no | yes | 10 | no | sand | normal | many | 12 | 12 |
| CUE_S3_S_F_C10 | CUE | SUP | Forest | 40.475659 | -2.457285 | 793 | moderate | ne | no | tree | no | yes | 10 | no | sand | normal | many | 12 | 12 |
| CUE_S4_D_F_D7 | CUE | DEEP | Forest | 40.486657 | -2.456571 | 853 | moderate | n | no | free_soil | no | yes | 5 | no | sand | normal | normal | 15 | 15 |
| CUE_S4_S_F_D10 | CUE | SUP | Forest | 40.486657 | -2.456571 | 853 | moderate | n | no | free_soil | no | yes | 5 | no | sand | normal | normal | 15 | 15 |
| CUE_S5_D_G_E7 | CUE | DEEP | Grassland | 40.485739 | -2.468131 | 817 | high | nw | yes | free_soil | no | yes | 0 | few | sand | normal | many | 12.5 | 20 |
| CUE_S5_S_G_E10 | CUE | SUP | Grassland | 40.485739 | -2.468131 | 817 | high | nw | yes | free_soil | no | yes | 0 | few | sand | normal | many | 12.5 | 20 |
| CUE_S6_D_G_F7 | CUE | DEEP | Grassland | 40.489145 | -2.470439 | 818 | moderate | s | yes | free_soil | no | yes | 0 | no | sand | normal | many | 12 | 18 |
| CUE_S6_S_G_F10 | CUE | SUP | Grassland | 40.489145 | -2.470439 | 818 | moderate | s | yes | free_soil | no | yes | 0 | no | sand | normal | many | 12 | 18 |
| CUE_S7_D_G_G7 | CUE | DEEP | Grassland | 40.5033 | -2.459098 | 873 | moderate | se | yes | free_soil | no | yes | 0 | few | sand | normal | many | 14 | 15.5 |
| CUE_S7_S_G_G10 | CUE | SUP | Grassland | 40.5033 | -2.459098 | 873 | moderate | se | yes | free_soil | no | yes | 0 | few | sand | normal | many | 14 | 15.5 |
| CUE_S8_D_F_H7 | CUE | DEEP | Forest | 40.480068 | -2.449797 | 851 | moderate | n | no | batter | no | yes | 3 | few | sand | high | many | 13 | 20 |
| CUE_S8_S_F_H10 | CUE | SUP | Forest | 40.480068 | -2.449797 | 851 | moderate | n | no | batter | no | yes | 3 | few | sand | high | many | 13 | 20 |
| CUE_S9_D_F_A8 | CUE | DEEP | Forest | 40.524092 | -2.449656 | 1129 | high | n | no | free_soil | few | yes | 10 | many | sand | normal | many | 11 | 8.8 |
| CUE_S9_S_F_A11 | CUE | SUP | Forest | 40.524092 | -2.449656 | 1129 | high | n | no | free_soil | few | yes | 10 | many | sand | normal | many | 11 | 8.8 |

| | | | | | | | | | | | | | | | | | | | |
|---|---|---|---|---|---|---|---|---|---|---|---|---|---|---|---|---|---|---|---|
| GRA_S1_D_F_A1 | GRA | DEEP | Forest | 36.748326 | -5.490475 | 706 | moderate | w | no | tree | few | yes | 3 | few | sand | low | few | 14 | 23.3 |
| GRA_S1_S_F_A4 | GRA | SUP | Forest | 36.748326 | -5.490475 | 706 | moderate | w | no | tree | few | yes | 3 | few | sand | low | few | 14 | 23.3 |
| GRA_S10_D_F_C2 | GRA | DEEP | Forest | 36.7543 | -5.443551 | 782 | no | nw | no | tree | few | yes | 5 | no | clay | high | many | 14.5 | 8 |
| GRA_S10_S_F_B5 | GRA | SUP | Forest | 36.7543 | -5.443551 | 782 | no | nw | no | tree | few | yes | 5 | no | clay | high | many | 14.5 | 8 |
| GRA_S11_D_F_D2 | GRA | DEEP | Forest | 36.756312 | -5.455879 | 633 | moderate | e | no | tree | no | yes | 3 | few | clay | normal | normal | 13 | 22 |
| GRA_S11_S_F_C5 | GRA | SUP | Forest | 36.756312 | -5.455879 | 633 | moderate | e | no | tree | no | yes | 3 | few | clay | normal | normal | 13 | 22 |
| GRA_S12_D_G_E2 | GRA | DEEP | Grassland | 36.790817 | -5.500449 | 400 | moderate | e | no | free_soil | no | yes | 0 | no | clay | low | few | 14 | 27 |
| GRA_S12_S_G_D5 | GRA | SUP | Grassland | 36.790817 | -5.500449 | 400 | moderate | e | no | free_soil | no | yes | 0 | no | clay | low | few | 14 | 27 |
| GRA_S13_D_G_F2 | GRA | DEEP | Grassland | 36.775142 | -5.50175 | 316 | moderate | w | yes | free_soil | no | yes | 0 | no | clay | low | normal | 14 | 16 |
| GRA_S13_S_G_G6 | GRA | SUP | Grassland | 36.775142 | -5.50175 | 316 | moderate | w | yes | free_soil | no | yes | 0 | no | clay | low | normal | 14 | 16 |
| GRA_S14_D_G_G2 | GRA | DEEP | Grassland | 36.772159 | -5.501121 | 304 | moderate | w | yes | free_soil | no | yes | 0 | no | clay | low | normal | 14 | 27 |
| GRA_S14_S_G_E5 | GRA | SUP | Grassland | 36.772159 | -5.501121 | 304 | moderate | w | yes | free_soil | no | yes | 0 | no | clay | low | normal | 14 | 27 |
| GRA_S15_D_F_H2 | GRA | DEEP | Forest | 36.755493 | -5.453507 | 660 | high | sw | no | stone | no | yes | 10 | many | clay | normal | normal | 12.5 | 10 |
| GRA_S15_S_F_G5 | GRA | SUP | Forest | 36.755493 | -5.453507 | 660 | high | sw | no | stone | no | yes | 10 | many | clay | normal | normal | 12.5 | 10 |
| GRA_S16_D_F_A3 | GRA | DEEP | Forest | 36.794996 | -5.394465 | 708 | high | nw | no | batter | no | yes | 3 | few | clay | low | normal | 12 | 18 |
| GRA_S16_S_F_F5 | GRA | SUP | Forest | 36.794996 | -5.394465 | 708 | high | nw | no | batter | no | yes | 3 | few | clay | low | normal | 12 | 18 |
| GRA_S17_D_F_B3 | GRA | DEEP | Forest | 36.78609 | -5.409673 | 771 | high | n | no | tree | few | yes | 3 | few | clay | low | few | 12 | 16.5 |
| GRA_S17_S_F_E6 | GRA | SUP | Forest | 36.78609 | -5.409673 | 771 | high | n | no | tree | few | yes | 3 | few | clay | low | few | 12 | 16.5 |
| GRA_S18_D_F_C3 | GRA | DEEP | Forest | 36.777118 | -5.402857 | 804 | high | n | yes | free_soil | no | yes | 3 | few | clay | low | few | 11 | 25 |
| GRA_S18_S_F_F6 | GRA | SUP | Forest | 36.777118 | -5.402857 | 804 | high | n | yes | free_soil | no | yes | 3 | few | clay | low | few | 11 | 25 |
| GRA_S19_D_G_B2 | GRA | DEEP | Grassland | 36.783147 | -5.411511 | 748 | no | nw | yes | batter | no | yes | 0 | few | clay | low | few | 12 | 25 |
| GRA_S19_S_G_H5 | GRA | SUP | Grassland | 36.783147 | -5.411511 | 748 | no | nw | yes | batter | no | yes | 0 | few | clay | low | few | 12 | 25 |
| GRA_S2_D_F_B1 | GRA | DEEP | Forest | 36.752437 | -5.489837 | 680 | high | w | no | batter | no | yes | 3 | no | sand | normal | normal | 13 | 27 |
| GRA_S2_S_F_B4 | GRA | SUP | Forest | 36.752437 | -5.489837 | 680 | high | w | no | batter | no | yes | 3 | no | sand | normal | normal | 13 | 27 |
| GRA_S20_D_F_D3 | GRA | DEEP | Forest | 36.790219 | -5.417923 | 894 | high | s | no | tree | no | yes | 5 | many | clay | normal | many | 13 | 15 |
| GRA_S20_S_F_C6 | GRA | SUP | Forest | 36.790219 | -5.417923 | 894 | high | s | no | tree | no | yes | 5 | many | clay | normal | many | 13 | 15 |
| GRA_S21_D_F_E3 | GRA | DEEP | Forest | 36.781208 | -5.427536 | 973 | moderate | ne | no | tree | no | yes | 10 | few | clay | low | few | 12 | 24 |
| GRA_S21_S_F_H6 | GRA | SUP | Forest | 36.781208 | -5.427536 | 973 | moderate | ne | no | tree | no | yes | 10 | few | clay | low | few | 12 | 24 |
| GRA_S22_D_G_G3 | GRA | DEEP | Grassland | 36.776266 | -5.435704 | 935 | moderate | w | yes | free_soil | few | yes | 0 | many | clay | normal | normal | 14 | 23 |
| GRA_S22_S_G_A6 | GRA | SUP | Grassland | 36.776266 | -5.435704 | 935 | moderate | w | yes | free_soil | few | yes | 0 | many | clay | normal | normal | 14 | 23 |
| GRA_S23_D_F_F3 | GRA | DEEP | Forest | 36.772708 | -5.440091 | 887 | high | n | no | tree | few | yes | 5 | few | clay | normal | few | 12 | 22 |
| GRA_S23_S_F_D6 | GRA | SUP | Forest | 36.772708 | -5.440091 | 887 | high | n | no | tree | few | yes | 5 | few | clay | normal | few | 12 | 22 |
| GRA_S24_D_G_H3 | GRA | DEEP | Grassland | 36.770455 | -5.457361 | 481 | moderate | s | no | free_soil | few | yes | 0 | many | clay | low | few | 14 | 20 |

| | | | | | | | | | | | | | | | | | | | |
|---|---|---|---|---|---|---|---|---|---|---|---|---|---|---|---|---|---|---|---|
| GRA_S24_S_G_B6 | GRA | SUP | Grassland | 36.770455 | -5.457361 | 481 | moderate | s | no | free_soil | few | yes | 0 | many | clay | low | few | 14 | 20 |
| GRA_S3_D_F_C1 | GRA | DEEP | Forest | 36.756077 | -5.493362 | 544 | moderate | w | no | batter | few | yes | 3 | many | clay | normal | many | 14 | 17 |
| GRA_S3_S_F_C4 | GRA | SUP | Forest | 36.756077 | -5.493362 | 544 | moderate | w | no | batter | few | yes | 3 | many | clay | normal | many | 14 | 17 |
| GRA_S4_D_G_D1 | GRA | DEEP | Grassland | 36.750688 | -5.433064 | 745 | moderate | sw | yes | batter | no | yes | 0 | no | clay | low | few | 14 | 45 |
| GRA_S4_S_G_D4 | GRA | SUP | Grassland | 36.750688 | -5.433064 | 745 | moderate | sw | yes | batter | no | yes | 0 | no | clay | low | few | 14 | 45 |
| GRA_S5_D_G_E1 | GRA | DEEP | Grassland | 36.750095 | -5.4272 | 743 | moderate | sw | yes | batter | no | yes | 0 | no | clay | low | normal | 14 | 46 |
| GRA_S5_S_G_E4 | GRA | SUP | Grassland | 36.750095 | -5.4272 | 743 | moderate | sw | yes | batter | no | yes | 0 | no | clay | low | normal | 14 | 46 |
| GRA_S6_D_G_F1 | GRA | DEEP | Grassland | 36.751254 | -5.433662 | 756 | moderate | w | yes | batter | no | yes | 0 | many | clay | low | normal | 15 | 23 |
| GRA_S6_S_G_F4 | GRA | SUP | Grassland | 36.751254 | -5.433662 | 756 | moderate | w | yes | batter | no | yes | 0 | many | clay | low | normal | 15 | 23 |
| GRA_S7_D_G_G1 | GRA | DEEP | Grassland | 36.766561 | -5.343489 | 638 | moderate | w | no | free_soil | few | yes | 0 | few | clay | normal | normal | 15 | 24 |
| GRA_S7_S_G_H4 | GRA | SUP | Grassland | 36.766561 | -5.343489 | 638 | moderate | w | no | free_soil | few | yes | 0 | few | clay | normal | normal | 15 | 24 |
| GRA_S8_D_G_H1 | GRA | DEEP | Grassland | 36.764764 | -5.346152 | 647 | moderate | n | yes | batter | no | yes | 0 | no | clay | normal | normal | 15 | 25 |
| GRA_S8_S_G_G4 | GRA | SUP | Grassland | 36.764764 | -5.346152 | 647 | moderate | n | yes | batter | no | yes | 0 | no | clay | normal | normal | 15 | 25 |
| GRA_S9_D_G_A2 | GRA | DEEP | Grassland | 36.758131 | -5.355284 | 688 | moderate | n | yes | free_soil | no | yes | 0 | few | clay | normal | few | 13 | 23 |
| GRA_S9_S_G_A5 | GRA | SUP | Grassland | 36.758131 | -5.355284 | 688 | moderate | n | yes | free_soil | no | yes | 0 | few | clay | normal | few | 13 | 23 |

**Table S2.** Metabarcoding primers used for the amplification of the bc3' fragment corresponding to 418 bp of the 3' end of the COI barcode region. PCRs were performed in 15 µL reaction volumes containing 3 mM $MgCl_2$, 0.2 mM dNTPs, 0.4 µM each primer and 0.5 U Taq DNA polymerase (TaKaRa Biosystems) and 1 µL of DNA bulk extraction (raw, 1/10, 1/100 dilutions). We used a cycling profile of 95ºC for 4 min; 28 cycles of 95ºC for 30 s; 48ºC for 30 s; 72ºC for 3 min, and a final extension of 72ºC for 10 min. Primers included a tail corresponding to the Illumina P5 and P7 sequencing adapters for subsequent library preparation following Nextera XT Index Kit; Illumina, San Diego, CA, USA.

| **Primer*** | | **Sequence (5'–3')** | **Reference** |
|---|---|---|---|
| Ill_B_F | **F** | CCIGAYATRGCITTYCCICG | Shokralla et a. 2015 |
| Fol-degen-rev | **R** | TANACYTCNGGRTGNCCRAARAAYCA | Yu et al. 2012 |

*F, forward; R, reverse.

**Table S3.** Differences in the mesofauna richness by sample (alpha diversity) for the different habitats and soil layers within each of the three local settings and at the multiple genetic similarity levels.

| | | haplotypes | | 1% lineages | | 2% lineages | | 3% lineages | | 4% lineages | | 5% lineages | | 6% lineages | | 8% lineages | |
|---|---|---|---|---|---|---|---|---|---|---|---|---|---|---|---|---|---|
| | | *F* | *p* | *F* | *p* | *F* | *p* | *F* | *p* | *F* | *p* | *F* | *p* | *F* | *p* | *F* | *p* |
| **GRA** | *habitat* | 7.80 | 0.011 | 9.67 | 0.005 | 10.47 | 0.004 | 11.33 | 0.003 | 11.92 | 0.002 | 11.69 | 0.002 | 11.24 | 0.003 | 10.90 | 0.003 |
| | *soil layer* | 38.16 | 0.000 | 29.53 | 0.000 | 22.79 | 0.000 | 18.55 | 0.000 | 17.46 | 0.000 | 16.92 | 0.000 | 17.39 | 0.000 | 17.02 | 0.000 |
| | *FOREST layer* | 32.00 | 0.000 | 22.66 | 0.001 | 16.56 | 0.002 | 14.08 | 0.003 | 13.76 | 0.003 | 13.31 | 0.004 | 14.00 | 0.003 | 13.03 | 0.004 |
| | *GRASSLAND layer* | 13.61 | 0.004 | 11.35 | 0.006 | 8.45 | 0.014 | 6.21 | 0.030 | 5.36 | 0.041 | 5.07 | 0.046 | 5.07 | 0.046 | 5.33 | 0.041 |
| **ALZ** | *habitat* | 0.59 | 0.451 | 0.24 | 0.629 | 0.11 | 0.749 | 0.07 | 0.794 | 0.01 | 0.935 | 0.09 | 0.762 | 0.02 | 0.886 | 0.01 | 0.920 |
| | *soil layer* | 10.31 | 0.004 | 10.26 | 0.004 | 9.93 | 0.004 | 9.47 | 0.005 | 8.12 | 0.009 | 7.29 | 0.013 | 7.55 | 0.012 | 8.50 | 0.008 |
| | *FOREST layer* | 6.85 | 0.024 | 8.67 | 0.013 | 9.18 | 0.012 | 8.87 | 0.013 | 8.67 | 0.013 | 7.86 | 0.017 | 8.21 | 0.015 | 8.33 | 0.015 |
| | *GRASSLAND layer* | 3.41 | 0.092 | 2.36 | 0.153 | 1.92 | 0.193 | 1.75 | 0.213 | 1.04 | 0.330 | 0.89 | 0.366 | 0.92 | 0.357 | 1.37 | 0.266 |
| **CUE** | *habitat* | 0.06 | 0.815 | 0.00 | 0.965 | 0.09 | 0.764 | 0.01 | 0.924 | 0.07 | 0.795 | 0.12 | 0.737 | 0.13 | 0.725 | 0.19 | 0.664 |
| | *soil layer* | 47.28 | 0.000 | 34.86 | 0.000 | 26.97 | 0.000 | 26.24 | 0.000 | 27.22 | 0.000 | 26.70 | 0.000 | 27.29 | 0.000 | 27.21 | 0.000 |
| | *FOREST layer* | 23.37 | 0.001 | 18.36 | 0.001 | 15.41 | 0.002 | 16.51 | 0.002 | 17.35 | 0.002 | 17.44 | 0.002 | 17.51 | 0.002 | 17.57 | 0.002 |
| | *GRASSLAND layer* | 22.20 | 0.001 | 15.42 | 0.002 | 10.97 | 0.007 | 9.59 | 0.010 | 9.85 | 0.009 | 9.42 | 0.011 | 9.80 | 0.010 | 9.71 | 0.010 |

**Table S4.** Differences in the compositional dissimilarity of mesofauna communities for the different habitats and soil layers within each of the three local settings and at the multiple genetic similarity levels.

| | | haplotypes | | | 1% lineages | | | 2% lineages | | | 3% lineages | | | 4% lineages | | | 5% lineages | | | 6% lineages | | | 8% lineages | | |
|---|---|---|---|---|---|---|---|---|---|---|---|---|---|---|---|---|---|---|---|---|---|---|---|---|---|
| | | *F* | $r^2$ | *p* | *F* | $r^2$ | *p* | *F* | $r^2$ | *p* | *F* | $r^2$ | *p* | *F* | $r^2$ | *p* | *F* | $r^2$ | *p* | *F* | $r^2$ | *p* | *F* | $r^2$ | *p* |
| **GRA** | *habitat* | 5.11 | 0.10 | 0.001 | 7.94 | 0.14 | 0.001 | 9.52 | 0.17 | 0.001 | 9.98 | 0.17 | 0.001 | 10.23 | 0.18 | 0.001 | 10.44 | 0.18 | 0.001 | 10.70 | 0.18 | 0.001 | 11.06 | 0.19 | 0.001 |
| | *soil layer* | 1.71 | 0.03 | 0.001 | 2.41 | 0.04 | 0.001 | 2.93 | 0.05 | 0.001 | 3.16 | 0.05 | 0.001 | 3.22 | 0.06 | 0.001 | 3.31 | 0.06 | 0.001 | 3.42 | 0.06 | 0.001 | 3.59 | 0.06 | 0.001 |
| **ALZ** | *habitat* | 4.20 | 0.09 | 0.001 | 5.63 | 0.11 | 0.001 | 6.55 | 0.12 | 0.001 | 6.67 | 0.13 | 0.001 | 6.84 | 0.13 | 0.001 | 7.09 | 0.13 | 0.001 | 7.21 | 0.13 | 0.001 | 7.36 | 0.13 | 0.001 |
| | *soil layer* | 1.53 | 0.03 | 0.001 | 1.92 | 0.04 | 0.001 | 2.10 | 0.04 | 0.001 | 2.49 | 0.05 | 0.001 | 2.61 | 0.05 | 0.001 | 2.78 | 0.05 | 0.001 | 2.80 | 0.05 | 0.001 | 3.06 | 0.06 | 0.001 |
| **CUE** | *habitat* | 4.74 | 0.09 | 0.001 | 6.08 | 0.11 | 0.001 | 6.02 | 0.11 | 0.001 | 6.34 | 0.12 | 0.001 | 5.80 | 0.11 | 0.001 | 5.95 | 0.11 | 0.001 | 5.97 | 0.11 | 0.001 | 6.06 | 0.11 | 0.001 |
| | *soil layer* | 2.05 | 0.04 | 0.001 | 2.59 | 0.05 | 0.001 | 3.35 | 0.06 | 0.001 | 3.55 | 0.06 | 0.001 | 3.84 | 0.07 | 0.001 | 3.86 | 0.07 | 0.001 | 3.86 | 0.07 | 0.001 | 3.92 | 0.07 | 0.001 |

**Table S5.** Correlations between the compositional dissimilarity matrixes of Acari, Collembola and Coleoptera communities within each of the three local settings and at the multiple genetic similarity levels.

| | | **haplotypes** | | | **1% lineages** | | | **2% lineages** | | | **3% lineages** | | | **4% lineages** | | | **5% lineages** | | | **6% lineages** | | | **8% lineages** | | |
|---|---|---|---|---|---|---|---|---|---|---|---|---|---|---|---|---|---|---|---|---|---|---|---|---|---|
| | | *F* | $r^2$ | *p* | *F* | $r^2$ | *p* | *F* | $r^2$ | *p* | *F* | $r^2$ | *p* | *F* | $r^2$ | *p* | *F* | $r^2$ | *p* | *F* | $r^2$ | *p* | *F* | $r^2$ | *p* |
| **GRA** | *Acari-Collembola* | 268.97 | 0.19 | 0.001 | 308.20 | 0.21 | 0.001 | 351.58 | 0.24 | 0.001 | 336.55 | 0.23 | 0.001 | 343.86 | 0.23 | 0.001 | 342.26 | 0.23 | 0.001 | 343.94 | 0.23 | 0.001 | 351.88 | 0.24 | 0.001 |
| | *Acari-Coleoptera* | 123.79 | 0.10 | 0.001 | 234.43 | 0.17 | 0.001 | 265.43 | 0.19 | 0.001 | 296.42 | 0.21 | 0.001 | 300.46 | 0.21 | 0.001 | 298.71 | 0.21 | 0.001 | 302.46 | 0.21 | 0.001 | 284.43 | 0.20 | 0.001 |
| | *Collembola-Coleoptera* | 79.09 | 0.07 | 0.001 | 130.06 | 0.10 | 0.001 | 159.03 | 0.12 | 0.001 | 153.55 | 0.12 | 0.001 | 149.55 | 0.12 | 0.001 | 163.74 | 0.13 | 0.001 | 184.30 | 0.14 | 0.001 | 181.63 | 0.14 | 0.001 |
| **ALZ** | *Acari-Collembola* | 199.63 | 0.16 | 0.001 | 159.97 | 0.13 | 0.001 | 119.35 | 0.10 | 0.001 | 118.83 | 0.10 | 0.001 | 136.51 | 0.12 | 0.001 | 154.93 | 0.13 | 0.001 | 144.78 | 0.12 | 0.001 | 133.02 | 0.11 | 0.001 |
| | *Acari-Coleoptera* | 98.30 | 0.08 | 0.001 | 152.21 | 0.12 | 0.001 | 160.76 | 0.12 | 0.001 | 159.34 | 0.12 | 0.001 | 167.38 | 0.13 | 0.001 | 174.57 | 0.13 | 0.001 | 178.83 | 0.14 | 0.001 | 189.17 | 0.14 | 0.001 |
| | *Collembola-Coleoptera* | 37.96 | 0.04 | 0.001 | 56.43 | 0.05 | 0.001 | 65.15 | 0.06 | 0.001 | 61.15 | 0.06 | 0.001 | 77.77 | 0.07 | 0.001 | 80.41 | 0.07 | 0.001 | 91.71 | 0.08 | 0.001 | 91.40 | 0.08 | 0.001 |
| **CUE** | *Acari-Collembola* | 141.72 | 0.11 | 0.001 | 108.64 | 0.09 | 0.001 | 88.05 | 0.07 | 0.001 | 99.44 | 0.08 | 0.001 | 82.01 | 0.07 | 0.001 | 72.85 | 0.06 | 0.001 | 80.06 | 0.07 | 0.001 | 75.33 | 0.06 | 0.001 |
| | *Acari-Coleoptera* | 121.03 | 0.10 | 0.001 | 206.77 | 0.16 | 0.001 | 185.29 | 0.14 | 0.001 | 188.04 | 0.14 | 0.001 | 174.93 | 0.13 | 0.001 | 151.16 | 0.12 | 0.001 | 153.06 | 0.12 | 0.001 | 159.01 | 0.12 | 0.001 |
| | *Collembola-Coleoptera* | 79.28 | 0.07 | 0.001 | 72.41 | 0.06 | 0.001 | 51.71 | 0.04 | 0.001 | 45.56 | 0.04 | 0.001 | 43.75 | 0.04 | 0.001 | 41.91 | 0.04 | 0.001 | 42.28 | 0.04 | 0.001 | 38.25 | 0.03 | 0.001 |

**Table S6.** Relationship between the compositional dissimilarity of mesofauna communities and the spatial distance (distance decay curves, $y= a*e^{-bx}$) within each of the three local settings and at the multiple genetic similarity levels.

| | | | | | | | | | | |
|---|---|---|---|---|---|---|---|---|---|---|
| **GRA** | **FOREST** | *Pseudo-$r^2$* | *intercept (a)* | *slope (b)* | *p* | **GRASSLAND** | *Pseudo-$r^2$* | *intercept (a)* | *slope (b)* | *p* |
| | **haplotypes** | 0.31 | 0.23 | -0.103 | 0.010 | **haplotypes** | 0.34 | 0.19 | -0.108 | 0.010 |
| | **1% lineages** | 0.19 | 0.33 | -0.060 | 0.010 | **1% lineages** | 0.23 | 0.23 | -0.064 | 0.010 |
| | **2% lineages** | 0.15 | 0.39 | -0.050 | 0.010 | **2% lineages** | 0.18 | 0.25 | -0.048 | 0.010 |
| | **3% lineages** | 0.13 | 0.41 | -0.042 | 0.010 | **3% lineages** | 0.17 | 0.26 | -0.046 | 0.010 |
| | **4% lineages** | 0.13 | 0.42 | -0.041 | 0.010 | **4% lineages** | 0.15 | 0.26 | -0.043 | 0.010 |
| | **5% lineages** | 0.13 | 0.42 | -0.039 | 0.010 | **5% lineages** | 0.15 | 0.27 | -0.041 | 0.010 |
| | **6% lineages** | 0.14 | 0.44 | -0.041 | 0.010 | **6% lineages** | 0.12 | 0.27 | -0.037 | 0.010 |
| | **8% lineages** | 0.12 | 0.44 | -0.036 | 0.010 | **8% lineages** | 0.10 | 0.27 | -0.032 | 0.010 |
| **ALZ** | **FOREST** | *Pseudo-$r^2$* | *intercept (a)* | *slope (b)* | *p* | **GRASSLAND** | *Pseudo-$r^2$* | *intercept (a)* | *slope (b)* | *p* |
| | **haplotypes** | 0.28 | 0.24 | -0.064 | 0.010 | **haplotypes** | 0.39 | 0.29 | -0.097 | 0.010 |
| | **1% lineages** | 0.30 | 0.33 | -0.057 | 0.010 | **1% lineages** | 0.36 | 0.34 | -0.086 | 0.010 |
| | **2% lineages** | 0.25 | 0.37 | -0.046 | 0.010 | **2% lineages** | 0.34 | 0.35 | -0.071 | 0.010 |
| | **3% lineages** | 0.21 | 0.38 | -0.039 | 0.010 | **3% lineages** | 0.24 | 0.35 | -0.054 | 0.010 |
| | **4% lineages** | 0.21 | 0.40 | -0.039 | 0.010 | **4% lineages** | 0.24 | 0.36 | -0.053 | 0.010 |
| | **5% lineages** | 0.23 | 0.43 | -0.041 | 0.010 | **5% lineages** | 0.20 | 0.37 | -0.048 | 0.010 |
| | **6% lineages** | 0.21 | 0.43 | -0.038 | 0.010 | **6% lineages** | 0.20 | 0.37 | -0.049 | 0.010 |
| | **8% lineages** | 0.21 | 0.44 | -0.037 | 0.010 | **8% lineages** | 0.20 | 0.38 | -0.048 | 0.010 |
| **CUE** | **FOREST** | *Pseudo-$r^2$* | *intercept (a)* | *slope (b)* | *p* | **GRASSLAND** | *Pseudo-$r^2$* | *intercept (a)* | *slope (b)* | *p* |
| | **haplotypes** | 0.36 | 0.30 | -0.100 | 0.010 | **haplotypes** | 0.14 | 0.24 | -0.061 | 0.010 |
| | **1% lineages** | 0.32 | 0.38 | -0.078 | 0.010 | **1% lineages** | 0.08 | 0.28 | -0.035 | 0.020 |
| | **2% lineages** | 0.31 | 0.42 | -0.072 | 0.010 | **2% lineages** | 0.07 | 0.29 | -0.029 | 0.020 |
| | **3% lineages** | 0.31 | 0.44 | -0.071 | 0.010 | **3% lineages** | 0.07 | 0.31 | -0.031 | 0.010 |
| | **4% lineages** | 0.30 | 0.45 | -0.070 | 0.010 | **4% lineages** | 0.09 | 0.31 | -0.034 | 0.020 |
| | **5% lineages** | 0.30 | 0.45 | -0.068 | 0.010 | **5% lineages** | 0.07 | 0.32 | -0.030 | 0.030 |
| | **6% lineages** | 0.30 | 0.45 | -0.069 | 0.010 | **6% lineages** | 0.06 | 0.32 | -0.028 | 0.050 |
| | **8% lineages** | 0.30 | 0.46 | -0.068 | 0.010 | **8% lineages** | 0.06 | 0.33 | -0.027 | 0.050 |

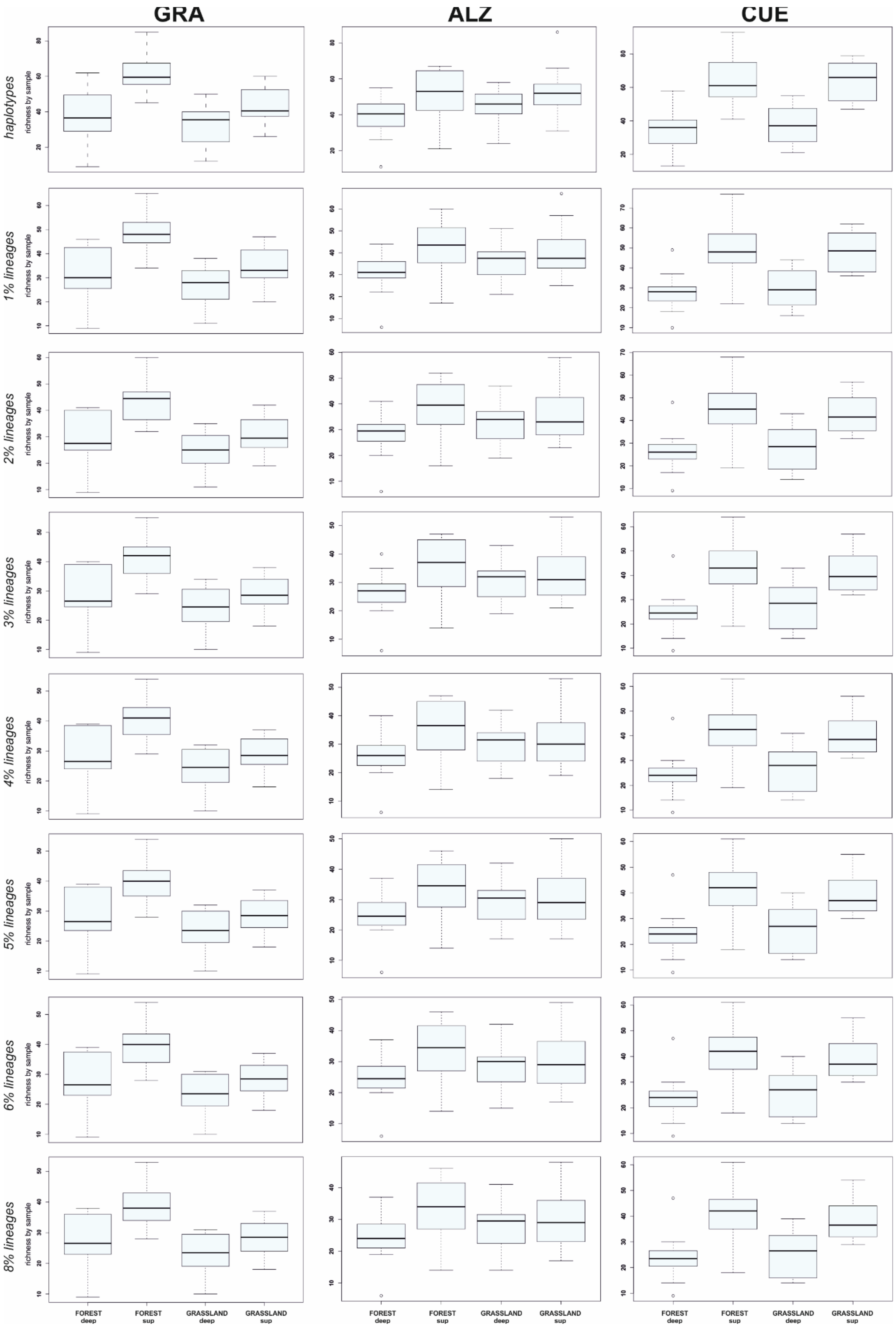

**Figure S1.** Mesofauna richness by sample (alpha diversity) for the different habitats and soil layers within each of the three local settings and at the multiple genetic similarity levels.

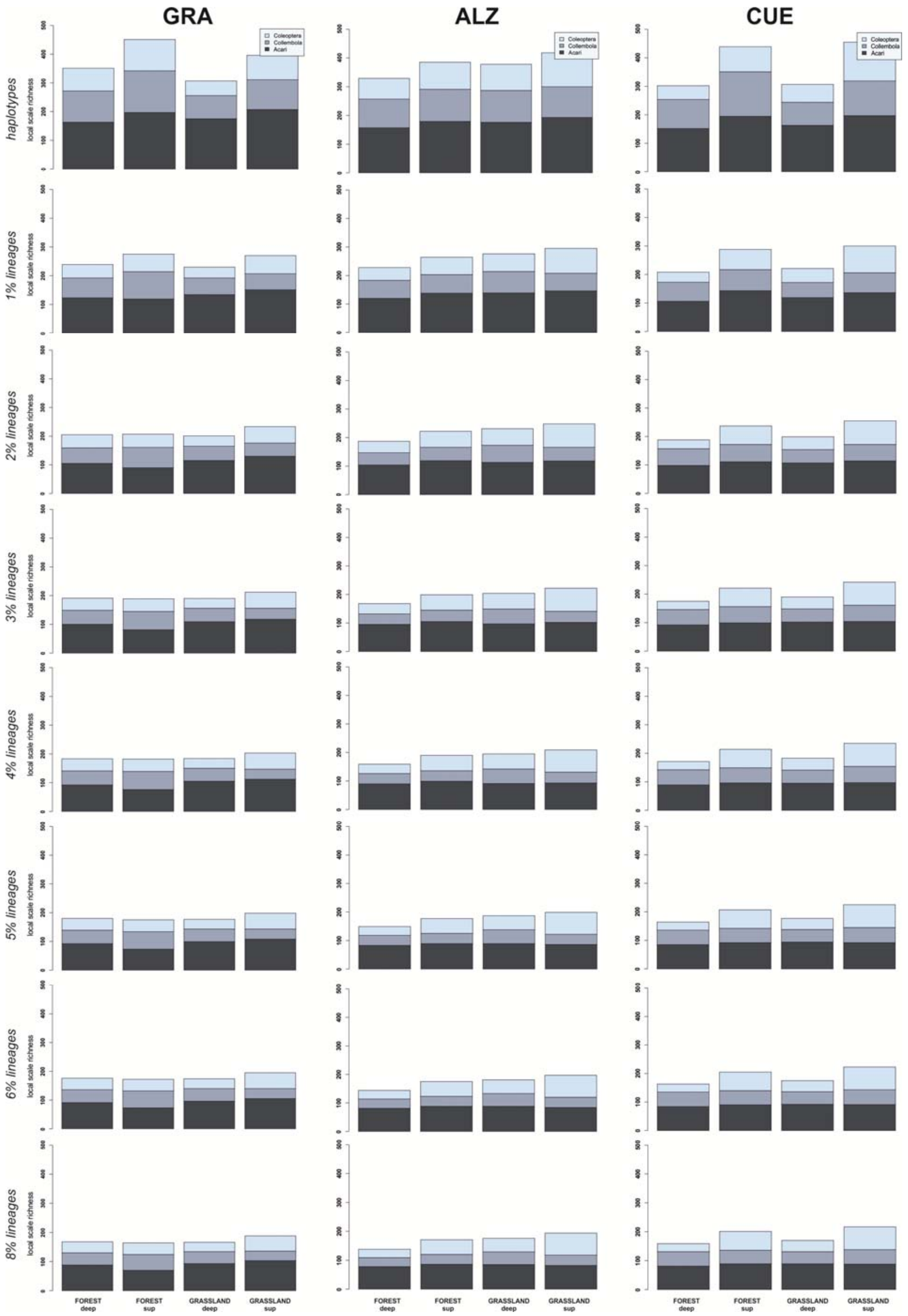

**Figure S2.** Mesofauna total accumulated richness (local scale richness) for the different habitats and soil layers within each of the three local settings and at the multiple genetic similarity levels.

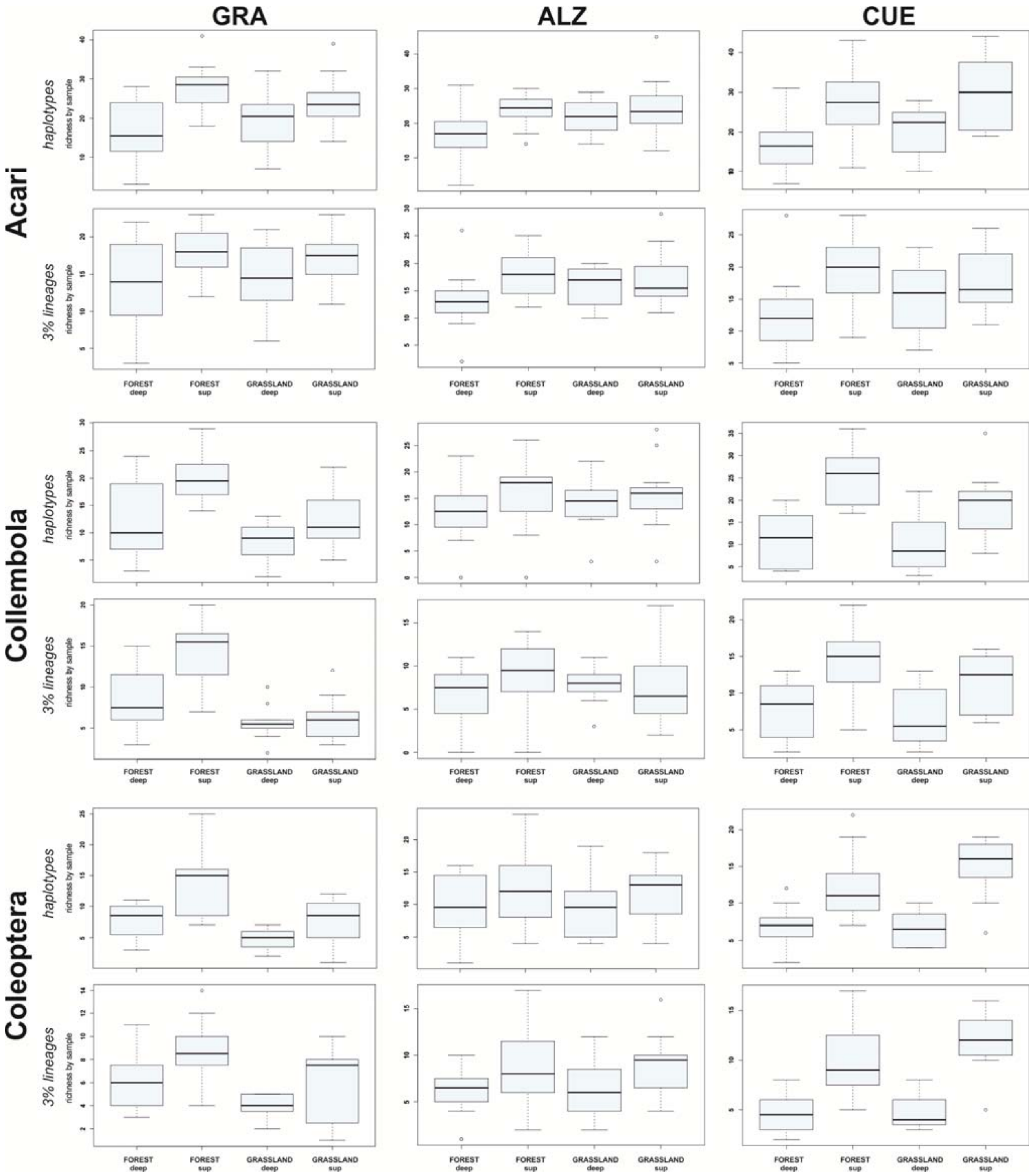

**Figure S3.** Acari, Collembola and Coleoptera richness by sample (alpha diversity) for the different habitats and soil layers within each of the three local settings and at the haplotype and 3% similarity levels.

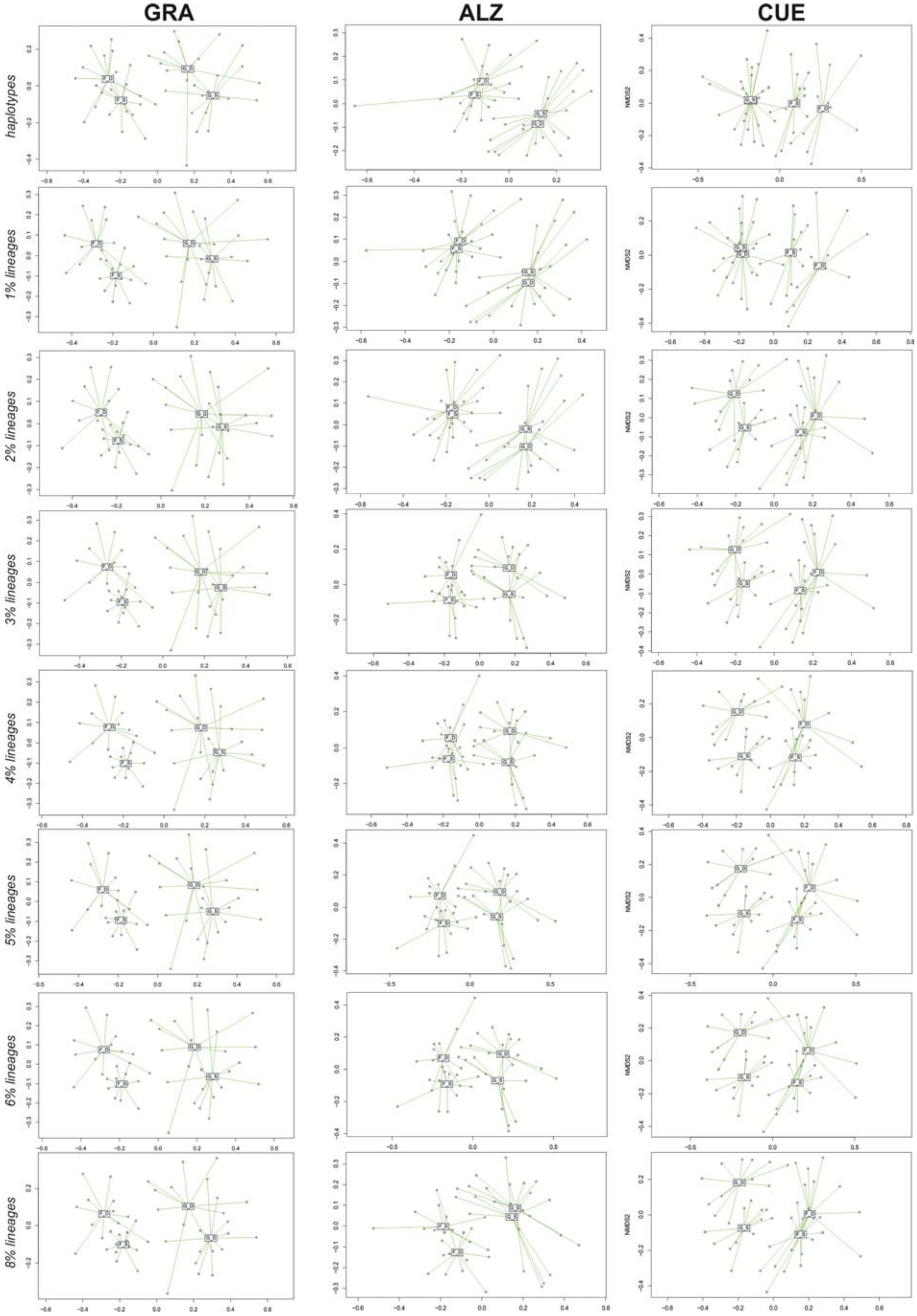

**Figure S4.** NMDS ordinations of the soil mesofauna samples according to the variation in community composition (Simpson index, βsim) for the different habitats and soil layers within each of the three local settings and at the multiple genetic similarity levels.
